# Effect of Mushroom-Bacteria Co-culture on Mushroom Growth and Antimicrobial Properties

**DOI:** 10.64898/2026.08.30.747672

**Authors:** Eunice Wang, Nicole Cavanaugh, Yinghao He, Yunrong Chai

**Affiliations:** Winchester High School, Winchester, MA; Biology Department, Northeastern University, Boston, MA

**Keywords:** Edible mushrooms, bacteria, fungi, co-culture, antimicrobials, shiitake mushroom

## Abstract

Edible mushrooms have been reported to have antimicrobial properties and other health benefits. This study aims to test the antimicrobial activities of several edible mushrooms from markets and test if co-culturing them with bacteria could induce stronger anti-bacterial properties. Commercial mushrooms, *Hericium erinaceus* (lion’s mane), *Pleurotus ostreatus* (oyster mushroom), *Lentinula edodes* (Shiitake) and *Agaricus bisporus* (button mushroom), were grown from strictly controlled/sterile substrates. Ethanol and water extracts from the mushrooms were prepared and tested against the bacteria *Escherichia coli*, *Pseudomonas aeruginosa*, *Staphylococcus aureus*, and *Bacillus subtilis*, and the fungus *Candida albicans* for antimicrobial activities. Shiitake water extract (SWE) showed strong antibacterial effects against all tested bacterial species, inhibitory effects on their biofilms, and antifungal activity. The antimicrobials in SWE seem to damage the cell wall and cell membrane of the bacteria, prefer weak acidic conditions, and are heat labile. Some antimicrobials are likely proteins and polysaccharides. In contrast, 3 other mushrooms displayed only weak antimicrobial effects. The fast-growing lion’s mane and oyster mushroom were co-cultured with different bacteria. The co-cultivation promoted the fruiting body development of lion’s mane. Co-culturing with *S. aureus* increased the anti-bacterial effects of lion’s mane against *S. aureus*, *E. coli* and particularly *B. subtilis*. Co-culturing the oyster mushroom with bacteria, especially *B. subtilis* and *P. aeruginosa*, boosted the mushroom’s growth. All tested bacteria, especially *S. aureus*, increased oyster mushroom’s anti-bacterial effect against *E. coli* and *B. subtilis*. The findings indicate that mushroom-bacteria co-culturing could have benefits both agriculturally and medicinally.

## Introduction

Edible mushrooms are not only consumed as food but also have been a sustainable source for bioactive compounds. Some mushrooms are commonly made into tinctures, which are concentrated liquid extracts created through a dual-extract of alcohol and water to get both the water-soluble and alcohol-soluble bioactive compounds, providing benefits such as boosted immunity, improved cognitive performance, anti-inflammatory effect, anti-cancer effect and antimicrobial effects (Valverde et al. 2015). Many antimicrobial agents have been obtained from various mushrooms (Alves et al. 2012; Anke et al. 1977). Antibiotic resistance is a critical global crisis, with drug resistance rising in over 40% of the pathogens monitored by the WHO (WHO. 2025). Most bacterial infections in humans involve biofilms, which are structured communities of microorganisms (bacteria, fungi, protists) embedded in a self-produced, slimy extracellular polymeric substance (EPS) matrix. Such structure makes biofilm-associated infections incredibly tough to clear because it protects the microbes from antimicrobials and makes microbes less responsive to antimicrobial treatments due to the stressful and competitive local environment within the biofilms. In contrast to the rapid rise of antimicrobial resistance, the development of new antimicrobials lags far behind with drug toxicity being one of the major hurdles. Edible mushrooms thus represent a novel, sustainable and relatively safe source of antimicrobial and anti-biofilm compounds (Valverde et al. 2015; Al Qutaibi et al. 2024).

In nature, mushrooms grow in competition with various bacteria and fungi which form biofilms in their local surroundings. However, commercial edible mushrooms are usually cultivated in much more controlled substrates, mostly from pasteurized or sterilized substrates. It would be interesting to examine the antimicrobial properties of the edible mushrooms from markets against bacteria and their biofilms, and if they can be induced to produce more antimicrobials when co-cultured with bacteria.

Button mushroom (*Agaricus bisporus*), shiitake mushroom (*Lentinula edodes*) and oyster mushroom (*Pleurotus ostreatus*) are the most widely cultivated mushrooms in the world, with *A. bisporus* accounting for roughly 90% of consumed mushrooms in the USA. Lion’s mane mushroom (*Hericium erinaceus*) is increasingly cultivated for both culinary and functional medicinal purposes. In this study, ethanol and water extracts were prepared from the fine powder of brown *A. bisporus*, *L. edodes*, *P. ostreatus var. columbinus* (blue oyster mushroom) and *H. erinaceus*, and tested for their antimicrobial effects on *Escherichia coli* MG1655 (Hayashi et al., 2006), *Pseudomonas aeruginosa* PAO1 (Stover et al., 2000), *Bacillus subtilis* NCIB 3610 (Nye et al., 2017) and *Staphylococcus aureus* HG003 (Sassi et al., 2014), respectively. Shiitake water extract (SWE) exhibited strong antimicrobial effects in the diffusion disk susceptibility test. It was thus further examined for its antibacterial efficiency against the four bacteria and their biofilms, its antifungal property, the stability of its antimicrobial property, and the possible compounds and mechanisms for the antimicrobial killing. Other mushrooms showed weaker antimicrobial effects from either their ethanol or water extracts. Two fast growing mushrooms, the blue oyster and lion’s mane, were chosen to perform the mushroom-bacteria co-culture experiments to see if co-culture can induce stronger antimicrobial effects in the mushrooms.

## Materials and Methods

### Materials

The brown button, shiitake, blue oyster and lion’s mane mushrooms were purchased from local markets for the initial antimicrobial property tests. Bacteria *E. coli* MG1655, *P. aeruginosa* PAO1, *S. aureus* HG003, *B. subtilis* NCIB 3610 were used for antimicrobial tests and mushroom-bacteria co-culture. Fungus *Candida albicans* SC5314 (Jones et al., 2004) was also used for antimicrobial testing. Sonifier Cell Disruptor (Model W185, Heats Systems-Ultrasonics, INC) was used to process mushroom powders to break cells for mushroom extracts. The Luria Bertani medium (1% w/v) was used for culturing bacteria, fungi and making LBA (1.5% w/v agar) plates. The Organic Blue Oyster Mushroom Grow Kits and Organic Lion’s Mane Mushroom Grow Kits (North Spore LLC, Portland, Maine), Terra Fungus mushroom grow tent, and Vivosun AeroStream H05 Intelligent Wi-Fi Humidifier were used for mushroom cultivation. Protease from *Aspergillus saitoi*, Type XIII (P2143-5G, Sigma-Aldrich, USA) and β-glucanase enzyme (LD Carlson company) were used for digesting proteins and the β-glucan in shiitake water extract, respectively. The dyes of SYTO™ 9 (green) and propidium iodide (PI, red) from the LIVE/DEAD® BacLightTM Bacterial Viability Kit were used for the cell envelope damage assay.

### Preparation of mushroom extracts by dual extraction method

200 g of each mushroom species were thoroughly cleaned, air dried overnight, transferred to the food dehydrator to dry at 35°C overnight until crispy and finally grounded into fine powder with a coffee grinder. One gram of fine mushroom powder was mixed thoroughly with 25 mL denatured 200-proof ethanol before being sonicated with ice bath by an ultrasonication cell disruptor for 30 rounds (1 round consisted of 30 seconds at level 4 followed by 1 minute rest).

The mixture was then centrifuged for 10 minutes at 10,000 rpm under 4°C and the supernatant was collected into a petri dish which was air dried overnight in a fume hood to get the dry ethanol extract powder. The solid sediment after centrifugation was mixed with 25 mL deionized water by vortex for 5 minutes, then centrifuged again at 10,000 rpm at 4°C. The supernatant was collected into a 50 mL conical tube which was deep frozen overnight at -80 °C before being lyophilized for 2 days to get the water extract dry powder. The ethanol extracts and water extracts were dissolved in dimethyl sulfoxide (DMSO) and water, respectively, with as high as possible concentration for stock before adjustment for experiments.

### Diffusion disk susceptibility test of mushroom extracts against bacteria

Bacterial lawn plates with *B. subtilis, E. coli, P. aeruginosa* and *S. aureus* were prepared by adding 200 µL of bacterial suspension at OD_600_=0.5 to each LBA plate, adding sterile glass beads, then shaking horizontally in different directions until the bacterial suspension was completely absorbed. Each plate was divided into 9 sections for the ethanol and water extract of four mushrooms and the negative control of DMSO, respectively. The paper diffusion disk of 0.7 cm loaded with 20 µL of the corresponding extract solution was placed on the center of the corresponding section. The experiment was set up in triplicates. All the plates were then incubated at 37°C for 18 ∼ 24 hours before they were taken out to record the diameter of clear zones.

### Minimum Inhibitory Concentration (MIC) and Minimum Bactericidal Concentration (MBC) Test

In a 96-well microtiter plate, each well of the first column was filled with 100 µL fresh LB broth, and each well of column 2 was filled with 100 µL of the starting concentration of SWE. Each well of column 3 to column 12 was filled with 50 µL LB broth. Starting from column 2, 50 µL of SWE was taken out and mixed with LB in column 3. This 2-fold dilution was repeated to column 11 where 50 µL of the final mixture was taken out and discarded. Overnight specific bacterial culture was diluted into LB broth at 1:1000 in a test tube, then 50µL of it was added to each well from column 2 to 12. In this way, column 1 was regarded as the negative growth control and column 12 as the growth control. The micrometer plate was then incubated at 37°C for 24 hours. From the first well without visible bacterial growth which was potentially the one with MIC to the next three wells with higher concentrations, all cultures were taken out respectively and centrifuged. The supernatant was discarded, and the remaining cell pellet was washed with PBS and resuspended in 100 µL PBS. Ten-fold serial dilution of the suspension was performed and plated in LBA plates and incubated overnight at 37°C for CFU counting. The concentration in the well with 10^6^ ∼ 10^7^ CFU/mL was taken as MIC and the well with 10^3^ ∼ 10^4^ CFU/mL was taken as MBC.

### Biofilm Inhibitory and Eradication Assay

For biofilm inhibitory assay, the 96-well micrometer plate was set up the same way as described in MIC assay. After a 24-hour incubation at 37°C, the liquid culture in each well was sucked out and discarded, and the well was washed with 100 µL PBS three times and then air dried. The crystal violet solution of 0.1% was added to the well to stain the biofilm for 30 min. The biofilm in the well was then washed with PBS three times again and air dried. The crystal violet in the biofilm was then dissolved by using 100 µL 95% ethanol before being placed into a spectrometer to read the absorbance at 595 nm. Because some wells had readings beyond the range, a 10-fold dilution of all wells was made into another 96-well plate to read with spectrometer again. In comparison to the negative and positive controls, the lowest concentration of SWE without biofilm growth was the minimum biofilm inhibitory concentration (MBIC). For the minimum biofilm eradication assay, LB broth was added as the negative control to column 1 of the 96-well plate. 100 µL bacterial culture diluted to OD_600_=0.05 from the overnight culture was added from column 2 to column 12. The plate was similarly incubated for 24 hours to have fully grown biofilm. The biofilm in each well was then washed with 100 µL PBS three times and remaining liquid in each well was carefully removed using a pipette. Two-fold serial dilution of SWE with the highest concentration of 100 mg/ mL was prepared in fresh LB solution and 100 µL of each was added to the corresponding biofilm wells in the 96-well plate, which was then incubated at 37°C for 24 hours. Similarly, each well was washed with PBS three times, air dried and stained with 0.1% crystal violet solution, washed by PBS and dissolved with 95% ethanol before reading the absorbance at 595 nm. The lowest concentration without biofilm growth was regarded as the minimum biofilm eradication concentration (MBEC).

### Killing curves of SWE against bacteria and *C. albicans*

The overnight culture of the bacteria and *C. albicans* were diluted at 1:20 into fresh LB medium in 96 well plates and incubated in the shaker at 37°C for 1∼1.5h before being treated with SWE of 50 mg/mL for *E. coli, P. aeruginosa*, 12.5 mg/mL for *B. subtilis* and 25 mg/mL for *S. aureus* and *C. albicans*. Samples were taken at t0, 0.5h, 1h and 2h-treatment for 10-fold serial dilution and LBA plating for CFU counting. The time points varied with different microbial species.

### Co-culturing lion’s mane and oyster mushroom with bacteria

Overnight cultures of *E. coli, P. aeruginosa, S. aureus and B. subtilis* were centrifuged to remove the media and then washed with PBS buffer three times. Then, the cells were resuspended into sterile deionized water and adjusted to the population density of OD_600_=1.0. All mushroom grow kits with substrate fully colonized with mycelium were labelled first with the mushroom bacteria combinations. The substrate blocks were taken out of the package. On the left side and right side, three evenly distributed spots were chosen as the injection points. For each mushroom-bacteria pair, 10 mL of the corresponding bacteria suspension was injected to the mushroom substrate package with a 10 mL syringe. The injections were performed in different angles and different depths in the substrate to ensure even coverage. The injection pores were wiped with ethanol before and after each injection before being sealed with a piece of tape sprayed with 70% ethanol. Then the co-culture substrate blocks were put back to the container box, making sure the orientation was the same, and incubated at around 20°C in dark for six days as the mycelium continued to grow. During the incubation, the kits were checked to see if the growth was still healthy. Then another round of injection of bacteria was performed in the same way, and the co-culture blocks were incubated overnight. The kit’s front side was cut open in an “X” shape to expose the substrate blocks to more air and promote fruiting. The blocks were then arranged on the shelves of the grow tent with a humidifier. The temperature was kept in the range of 15∼17°C, with 8 hours of light per day and a misting cycle of 1 hour of mist followed by 2 hours of rest. The daily growth was recorded. The lion’s mane mushroom was harvested when its teeth dropped downwards to around 1cm. The oyster mushroom was harvested when the edge of the fruiting body started to become flat or curled up. The mushroom was twisted off from the substrate, weighed, then torn into small stripes and air dried overnight before being transferred onto a dehydrator to further dry at 35**°**C for 10 hours. The crispy dry mushroom strips were ground to fine powder by a coffee grinder for 30 cycles, each cycle consisting of 5 seconds grinding and 5 seconds rest. The ethanol and water extracts were prepared using the mushroom powder and then subjected to diffusion disk susceptibility tests in triplicates as done before.

### Whole genome sequencing for the SWE resistant mutants of *B. subtilis*

Two *B. subtilis* mutant colonies (BM1 and BM2) grown in the clear zones of the diffusion disks in SWE were selected and purified by streaking the colonies on LBA plates. The mutant BM2 with an 8-fold increase in MIC was subjected to genomic DNA extraction followed by whole genome sequencing (Plasmidsaurus.com). The genome of the parent strain was sequenced as well as the reference. The assembled genome sequences (WT and BM2) were uploaded to NCBI and assigned with the BioSample accession numbers SAMN62780868 and SAMN62780869, respectively. Software NCBI nucleotide BLAST was used to align the genome sequence of the B. subtilis mutant BM2 with the wild type parent genome. The standalone executable version was downloaded from website (https://ftp.ncbi.nlm.nih.gov/blast/executables/blast+/LATEST/) to the local computer and the alignment analysis was conducted through command window.

### Genomic DNA extraction

Bacterial culture at OD_600_=0.5 in 2 mL was centrifuged at 8000 rpm (6800 x g) in a 2 mL Eppendorf tube for 2 minutes at room temperature. The cell pellet was re-suspended in 450 μL of autoclaved diH2O and had 50 μL of EDTA (0.5 M), 60 μL of lysozyme (stock at 20 mg/mL) and 60 μL of protease (P2143-5G. stock at 20 mg/mL) added to it. The tube was inverted 4-6 times, then incubated at 37°C for 1 hour before adding 650 μL of solution I (nuclei lysis solution) and inverting 4-6 times until the solution was clear. Then, 250 μL of solution II (protein precipitation solution) was added to the tube, which was vortexed for 20 seconds, left on ice for 10 minutes, then centrifuged at 14000 x g for 5 minutes. From the tube approximately 900 μL of the supernatant was transferred to a new tube where 600 μL of isopropanol was then added. The tube was inverted 4-6 times until a white precipitate was seen. The tube was left at room temperature for 3 minutes, then centrifuged for 5 minutes at 14000 x g. The supernatant was discarded. The pellet was washed by adding 600 μL of 70% ethanol and then centrifuging for 5 minutes at 14000 x g. The pellet was air dried and then re-suspended in 100 μL of autoclaved diH2O. The obtained DNA was stored at -20°C.

### Cell envelope damage assay

The LIVE/DEAD® BacLight^TM^ Bacterial Viability Kit (BacLight Kit) contains fluorescent dyes, SYTO 9 (green), and propidium iodide (PI) (red). Both dyes bind to nucleic acid. SYTO 9 is membrane permeable and can enter all cells, whereas PI is membrane impermeable and can only enter cells with compromised membranes, regarded as dead by the manual. However, cells killed by protein synthesis inhibition or DNA damage can still have intact cell envelopes, resulting in a green appearance under the microscope (Robertson et al. 2021). In this study, the assay was conducted to *B. subtilis* and *C. albicans.* The overnight culture was inoculated into fresh LB at 1:20 and incubated at 37°C for one hour, then placed to three wells of a 96-well plate with 200 μL per well, for the initial time point t0, 30 min and one hour of SWE treatment, with 12.5 mg/mL for *B. subtilis* and 25 mg/mL for *C. albicans*. At each time point, 10 μL samples were taken for a 10-fold serial dilution and LBA plating for CFU counting in triplicates. The rest of the cells were centrifuged, washed with PBS solution, resuspended in 5 μL of PBS, and had 1 μL of 10-fold dilution of the 2X stock solution from the LIVE/DEAD BacLight staining reagent mixture added. The solution was then incubated in the dark for 15 min, dropped on the slide in 3 μL and covered with the coverslip for observation under the Leica compound microscope with corresponding filter sets.

## Results

### SWE inhibits selected Gram-positive and -negative bacteria

The ethanol and water extracts of edible mushrooms were prepared as described in the Materials and Methods. Mushroom water extracts of 300 mg/mL and ethanol extracts of 150 mg/mL were used for the initial diffusion disk susceptibility test in triplicates against the Gram-negative bacteria *P. aeruginosa* and *E. coli* and the Gram-positive bacteria *S. aureus* and *B. subtilis.* The diameter of the filter paper diffusion disk is 0.7 cm. For shiitake mushroom, the water extract had a strong antimicrobial effect on all bacterial strains tested, but the ethanol extract showed minimal effect (Fig. 1). Button mushroom water extract had a minimal effect on *P. aeruginosa, E. coli and B. subtilis*, and its ethanol extract had some effect on *B. subtilis.* Both oyster mushroom water and ethanol extracts had a modest effect only on *B. subtilis*. No antimicrobial effect was observed from lion’s mane water extract, but a visible effect on *B. subtilis* was shown from its ethanol extract.

**Fig. 1.**
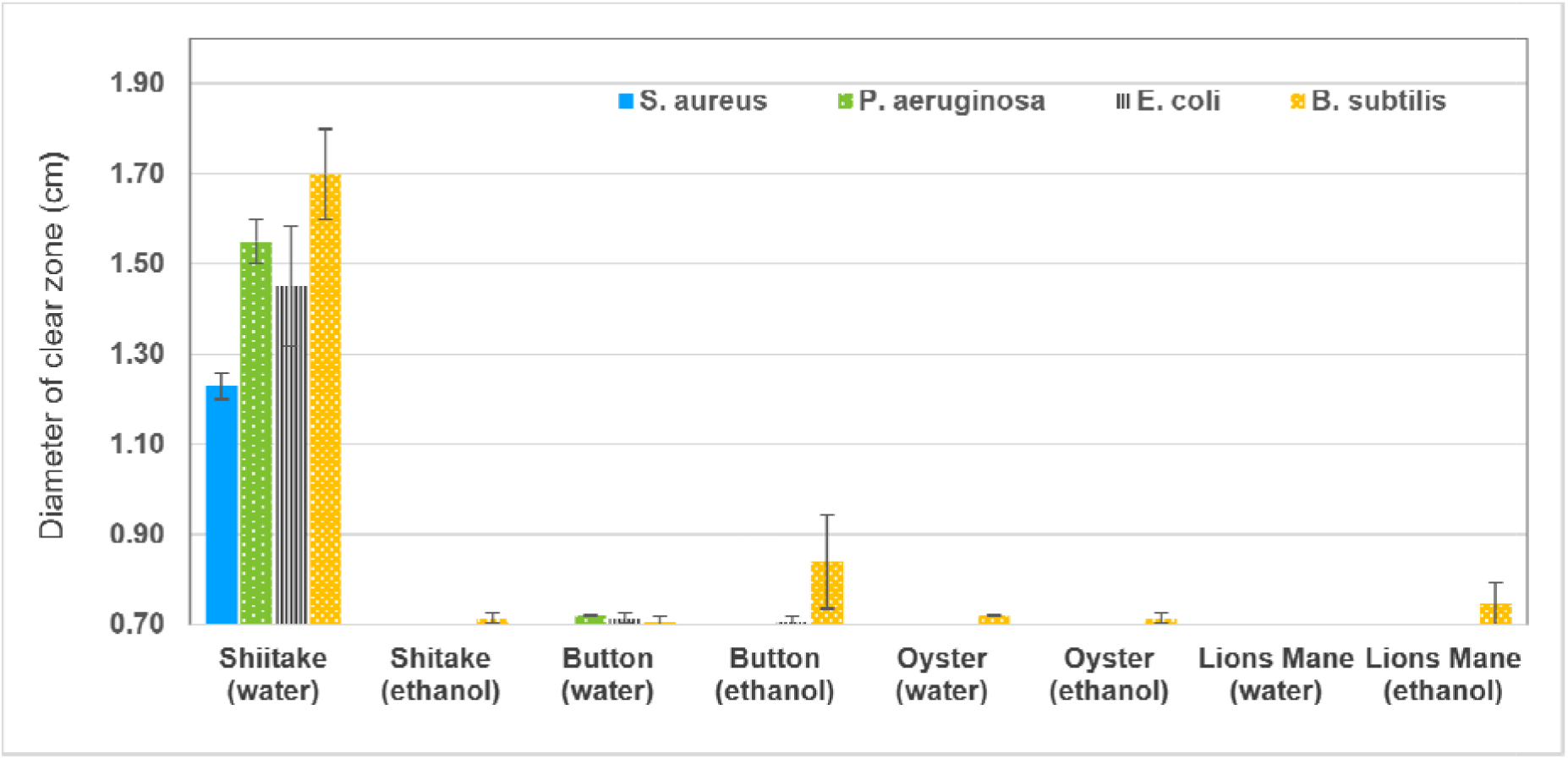
Diffusion disk susceptibility test of mushroom extracts on *S. aureus*, *P. aeruginosa*, *E. coli* and *B. subtilis.* Error bars: standard deviation (SD; n=3)

SWE showcased impressive broad antibacterial activities. Among the four tested bacteria, *E. coli* and *P. aeruginosa* are Gram-negative bacteria which have outer membrane to provide them with an extra layer of protection against antibiotics. *P. aeruginosa* is a well-known multidrug resistant opportunistic pathogen causing severe hospital acquired infections. *B. subtilis* and *S. aureus* are Gram-positive bacteria. For individuals with weakened immune systems, *S. aureus* can lead to severe conditions. Infections caused by some strains such as Methicillin-resistant *Staphylococcus aureus* (MRSA) are extremely hard to treat. Results in Fig. 1 show that all of them were efficiently inhibited by SWE in this test.

Based on test results in Fig. 1, the following studies were conducted in parallel in two directions: the antimicrobial properties of SWE, and the effect of mushroom-bacteria co-culture on the antimicrobial properties of mushrooms.

### Shiitake mushroom water extract inhibits the fungus *C. albicans*

*C. albicans* is an opportunistic pathogen causing oral infections or vaginal infection if the corresponding local microbiome of hosts is off balance. The diffusion disk susceptibility test of shiitake water and ethanol extract was performed on the lawns of *C. albicans* SC5314. The shiitake ethanol extract of 110 mg/mL showed no effect on the fungus, but the water extract of 100 mg/mL displayed a clear zone of 1.18 ± 0.02 cm (Supplementary Material Fig. S1).

### Determine MIC and MBC of SWE against bacteria and their biofilms

The MICs and MBCs of SWE to the four bacteria were tested on 96-well plates, and the result is shown in Fig. 2A. The Gram-positive bacteria, *S. aureus* and *B. subtilis* were more susceptible to the killing by SWE. This was expected since Gram-negative bacteria have an outer membrane which is an asymmetric lipid bilayer coated with lipopolysaccharides, inserted with porins, and usually parts of antibiotic efflux pumps, giving the bacteria additional mechanisms to fight against antibiotics.

**Fig. 2.**
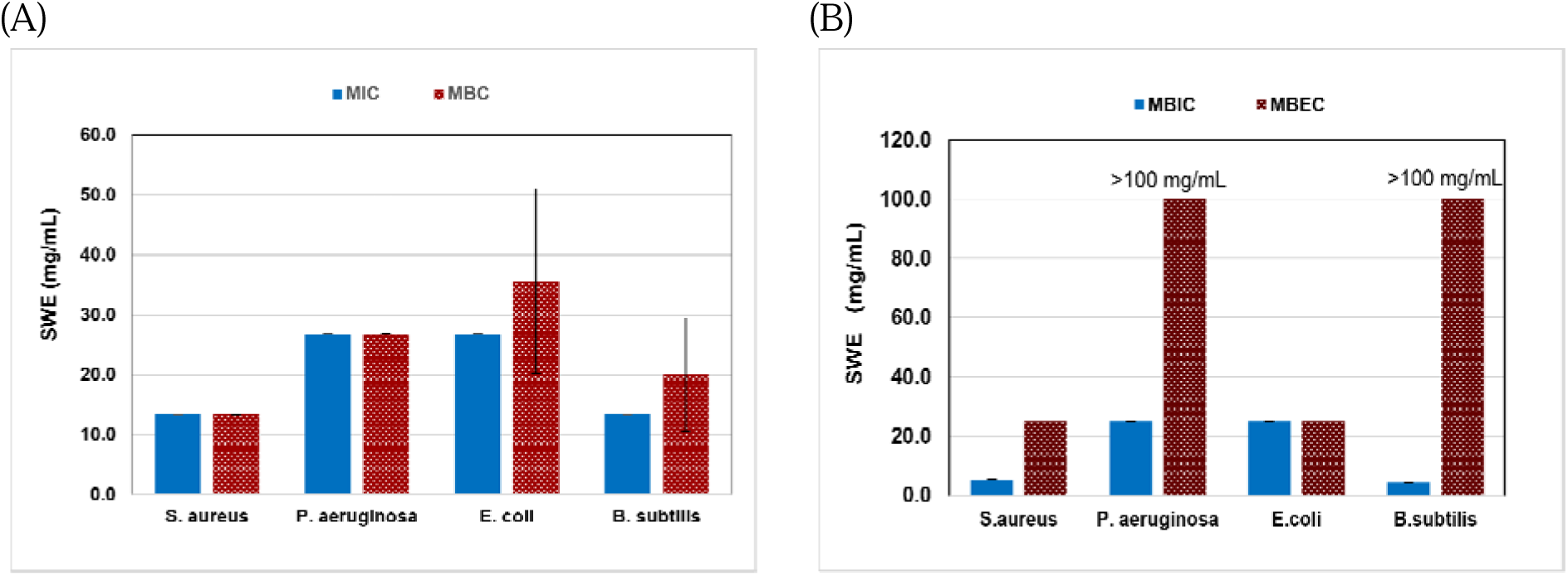
Susceptibility of *S. aureus, P. aeruginosa, E. coli* and *B. subtilis* to SWE (A) MIC and MBC to planktonic cells. (B) MBIC and MBEC by crystal violet staining method. Error bars: standard deviation (SD, n=3)

Biofilms are major culprits in hospital-acquired infections and frequently form on the surface of medical devices. Their structure provides protection to the bacterial or fungal cells within the biofilm from the antimicrobial treatments, making them more resistant than the free-living microbial cells. In this study, biofilm inhibition and eradication assays were performed with SWE by 2-fold serial dilution in flat-bottom 96-well plates as described in the methodology section. The MICs for biofilm inhibition in Fig. 2B showed that SWE is more effective at inhibiting the biofilm growth of gram-positive bacteria. Comparing Fig. 2A and Fig. 2B, the MICs for *B. subtilis* and *S. aureus* biofilm formation were relatively lower. This might be a result of the method used. The biofilm inhibitory and eradication assays were performed using the crystal violet staining method, which measured the biomass of biofilm instead of living cells.

Wells with living cells may not have biofilm formed, or loosely formed biofilm may be washed away during the procedures of the assay. The result of biofilm eradication assay clearly showed that biofilm is hard to eliminate. The highest concentration used could not eliminate the biofilm of *B. subtilis* and *P. aeruginosa*.

### SWE kills resting planktonic cells of bacteria and *C. albicans*

New SWE was prepared for the killing curve experiment, and it had a MIC of 3.13 mg/mL to *B. subtilis*. As shown in Fig. 3, for bacteria, the treatment started at the early stationary phase but SWE still achieved about 2-log killing to *E. coli*, 3-log to *B. subtilis* and remarkably 4-log to *P. aeruginosa* which is known to be naturally resistant to many antibiotics due to its low outer membrane permeability and active efflux pumps (Okamoto et al. 2001; Pang et al. 2019). At the start of SWE treatment, the population density of *S. aureus* was close to 10^10^ CFU/mL, but it still achieved a half log of killing. SWE wiped out *C. albicans* in 30 min (the detection limit is 10^4^ CFU/mL).

**Fig. 3.**
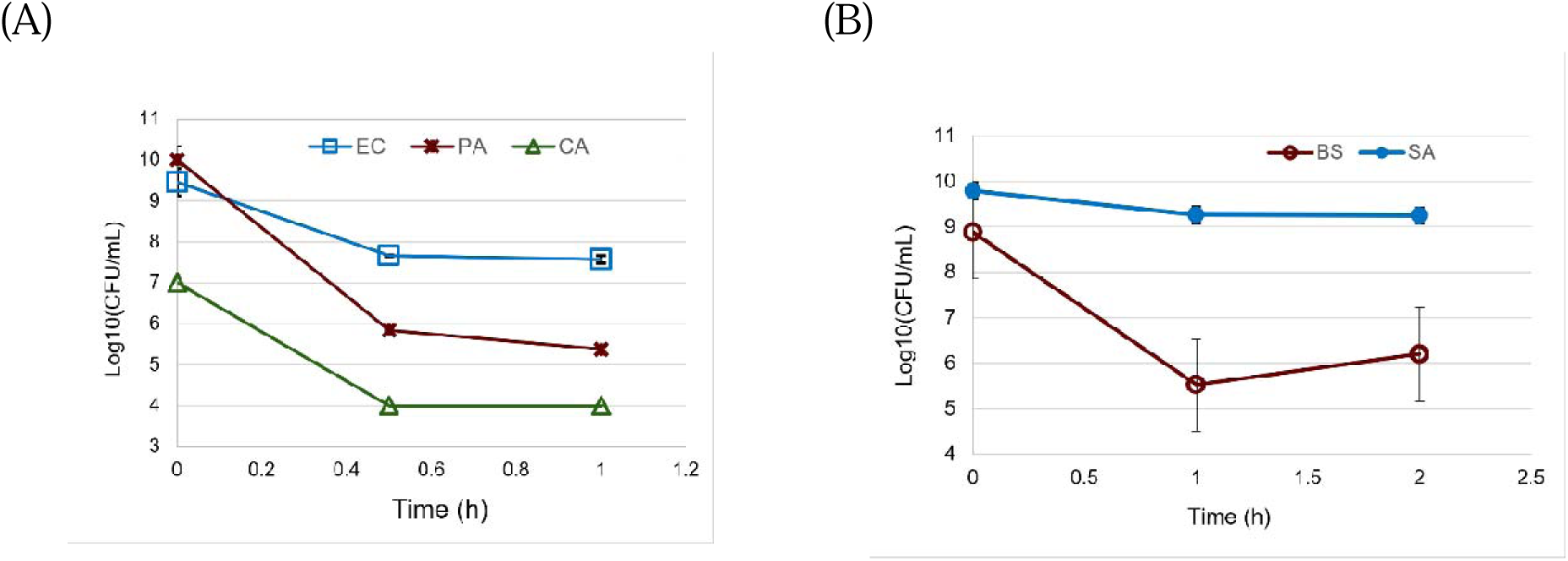
SWE time killing of (A) *E. coli* (EC), *P. aeruginosa* (PA) and *C. albicans* (CA). (B) *B. subtilis* (BS), *S. aureus* (SA). Error bars: standard deviation (SD; n=3)

### The antimicrobials in SWE are heat labile, favor weak acidic conditions and are sensitive to protease and β-glucanase

Aliquots of the SWE were incubated under room temperature (the control), 37°C, 43°C, 50°C, 70°C and 90°C for one hour, respectively, and then tested using the paper diffusion disks as shown in Fig.4 A. The antimicrobial activity did not have much change with temperature increasing from room temperature to 43°C. At 50°C, the size of the clear zone decreased, and the edge changed from fuzzy to sharp. The antimicrobial effect was completely lost at 70°C or above, indicating that the antimicrobial compounds in SWE were heat labile.

**Fig. 4.**
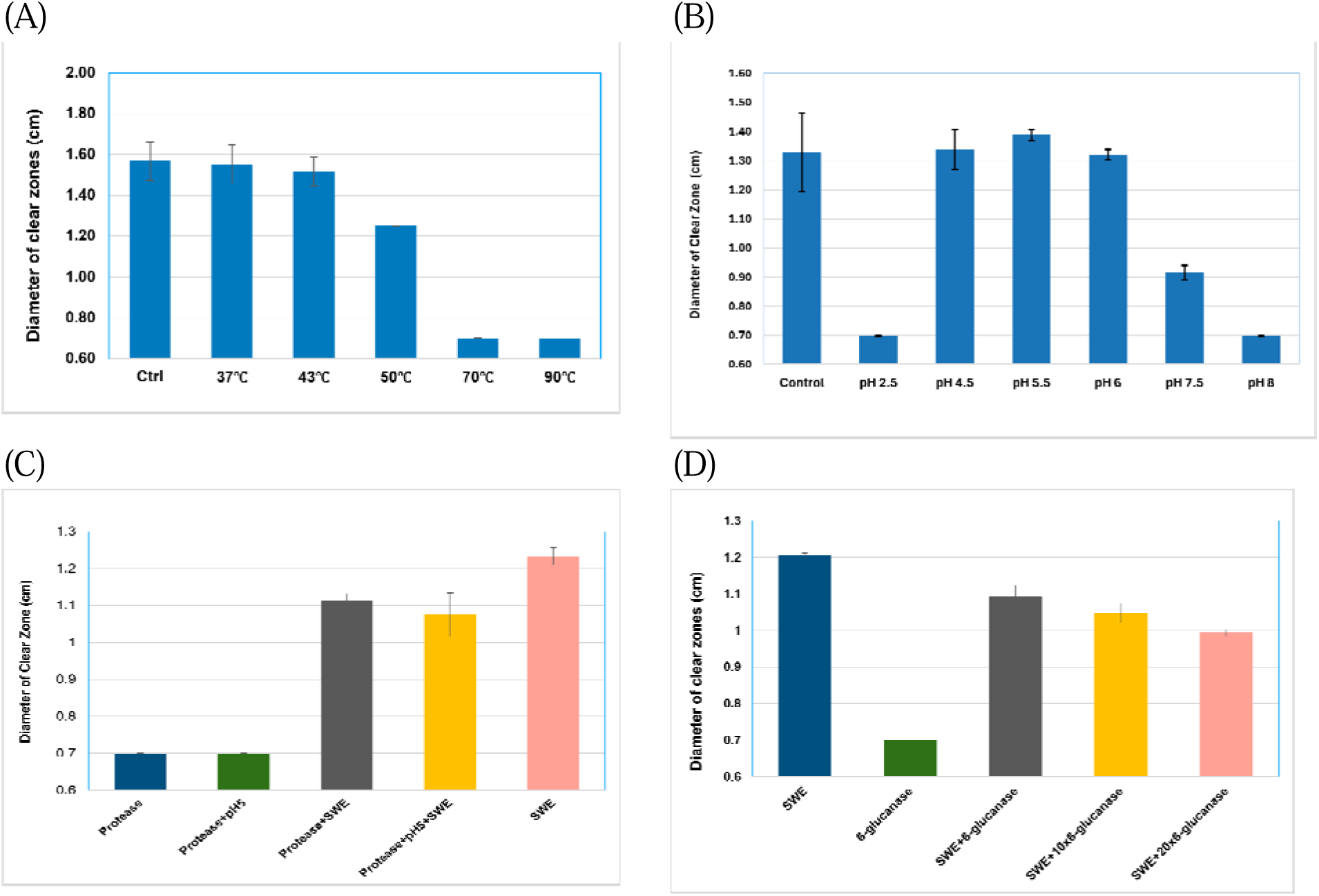
The stability of SWE antimicrobial property with (A) temperature; (B) pH; (C) protease digestion where the protease concentration was 2 mg/mL; and (D) β-glucanase digestion, where the glucanase concentration was 0.2 mg/mL, 2 mg/mL and 4mg/mL, respectively. The final concentration of SWE was 25 mg/mL. The diameter of the paper diffusion disk was 0.7 cm. Error bar: standard deviation (SD, n=3).

Fig. 4 B displayed the effect of pH on SWE antimicrobial property. To make SWE at different pH, the solutions of different pH were made in advance by diluting 6M HCl solution or 2 M NaOH solution with deionized water, then each solution was mixed with the SWE stock solution of 200 mg/mL, respectively, to form final solutions of 25 mg/mL SWE. The pH in Fig. 4B was the final pH of the mixture. The pH of SWE itself (control) was close to 6. SWE is a crude extract of shiitake mushroom, containing many molecular species which are weak acids or weak alkalines, behaving like a buffer. The test showed that SWE works better in weak acidic conditions. SWE at pH 2.5 did not generate a clear zone but made the lawn next to the diffusion disk thinner, showing a weak inhibiting effect (data not shown). The aqueous solutions of HCl or NaOH at different pH were tested with diffusion disks for their killing effect against *E. coli* (Supplementary Fig. S2). These aqueous solutions starting from pH 1.5 had no killing effect until the pH was increases to 12.5 which corresponded to 0.2 M NaOH, indicating that killing effect in Fig. 4B was completely from SWE antimicrobials.

To test if proteins or peptides contribute to the antimicrobial effects, SWE of 25 mg/mL was digested with the protease from *Aspergillus saitoi* (Sigma-Aldrich) at the final concentration of 2 mg/mL for one hour under 37°C before the susceptibility test with paper diffusion disk on the lawn of *E. coli*. The protease works better in an acidic condition. Therefore, parallel digestion samples were prepared in the aqueous solution of pH=5 as well. As shown in Fig. 4C, the protease digestion reduces the width of the clear ring by around 23% and protease treatment at pH=5 reduces the width by around 30%, indicating that SWE’s antimicrobial property comes partially from protein or peptides.

Polysaccharides have been reported to have antimicrobial effects through various mechanisms including disrupting membranes, inhibiting nuclear acid and protein synthesis, iron chelation, inhibiting biofilm formation, etc. (Liang et al. 2024). Shiitake mushrooms are rich in polysaccharides, and the most abundant and important is lentinan, a β-glucan. Lentinan has been reported to have an antimicrobial effect but was considered to work indirectly through its effect on immunity (Liu et al. 2019). We suspect that lentinan could be part of the antimicrobial activities. β-Glucanase enzyme (LD Carlson company) was used in 0.2 mg/mL (the recommended concentration by the manual), 2 mg/mL and 4 mg/mL, respectively, to digest SWE of 25 mg/mL under 37°C for one hour prior to the diffusion disk test. As shown in Fig. 4D, the digestion clearly decreased the width of the clear ring by around 20∼40%, and the decrease become dramatic with the increase of the enzyme concentration, indicating that lentinan may be able to directly kill *E. coli* or inhibits its growth.

### Antimicrobials in SWE damage bacterial cell envelope

A few *B. subtillis* colonies grew in the clear zones of the diffusion disk plates. They were selected and purified by streaking on fresh LBA plates. MIC assay of SWE to the mutants were then conducted to confirm their resistance. The mutant with 8 x MIC in comparison to the parent strain was selected for whole genome sequencing. The sequencing result confirmed a candidate list of genes with mutations (Supplementary Table S1) which was then analyzed preliminarily.

The mutated genes fall in a few categories: sporulation, quorum sensing, PBSX prophage, SPβ prophage and Toxin-antitoxin (TA) modules. The function of those candidate genes could be related to activation of sporulation, prophages, holins and hydrolases under environmental stresses such as treatment of the antimicrobials. The possible consequence is the loss of the integrity of cell envelope.

To examine if the antimicrobials in SWE damage cell envelope, the fluorescent dyes, SYTO 9 (green), and propidium iodide (PI) (red) from LIVE/DEAD BacLight Bacterial Viability Kit were used to stain *B. subtilis* and *C. albicans* cells before and after the treatment by SWE as described in the Material and Methods. The results were shown in Fig. 5A-D. The 1-h treatment made nearly a 3-log killing to *B. subtilis*; most cells shown in Fig. 5B were likely dead. A fraction of the cells were red, indicating that these cells did have compromised cell envelope because PI can only cross damaged cell membranes. A similar result was observed in *C. albicans* (Fig. 5C-D). Besides, the cells in Fig. 5C looked smoother and had a well-defined shape, whereas the cells in Fig. 5D appeared deformed and shrunken. To check if the microscope can capture cell lysis of bacterial and fungal cells, exponentially growing cells of *B. subtilis, S. aureus*, and *C. albicans* were mixed and treated with SWE at the concentration of 25 mg/mL, and 5 µL of the treated samples was loaded onto a slide for observation under microscope.

**Fig. 5.**
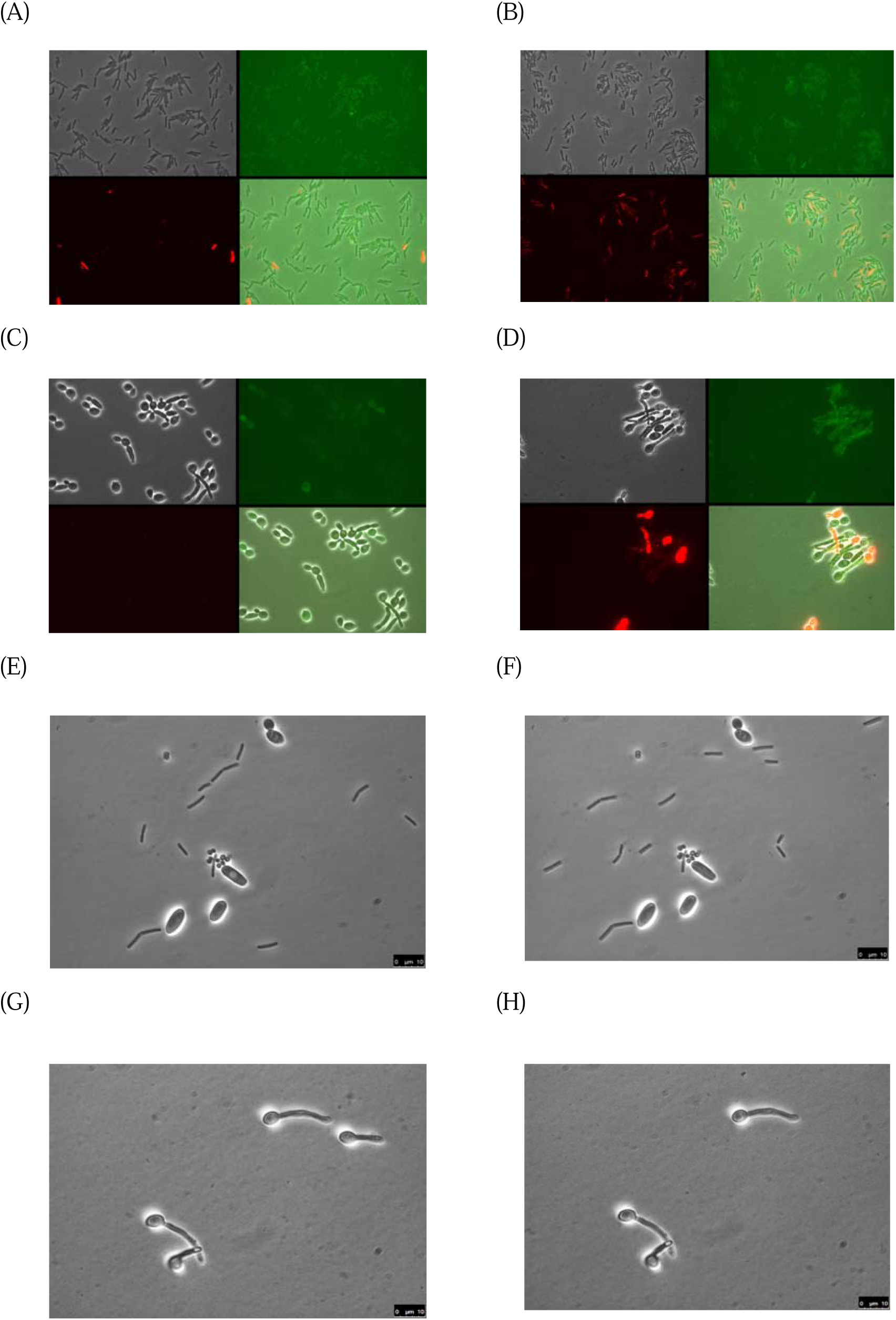
Cell envelope damage assays. (A) *B. subtilis* stained with SYTO 9 (green) and PI (red) before SWE treatment with a population density of 1.2 x10^8^ CFU/mL. (B) *B. subtilis* after 1-h treatment of SWE stained with SYTO 9 (green) and PI (red) with the number of viable cells being 2 x10^5^ CFU/mL. (C) *C. albicans* stained with SYTO 9 (green) and PI (red) before SWE treatment with a population density of 1x10^7^ CFU/mL. (D) *C. albicans* after 1-h treatment of SWE stained with SYTO 9 (green) and PI (red) with the number of viable cells being 1 x10^4^ CFU/mL. (E) *C. albicans, B. subtilis and S. aureus* cells with SWE treatment of around 40 min. A vacuole in *C. albicans* was clearly visible right before it collapsed. (F). The same view as in (E), right after the collapse of the vacuole in the *C. albicans* cell. (G) *C. albicans* in hyphal form, 7 min after SWE added onto the slide. (H) Same view as in (G) but 10 min after SWE was added onto the slide.

*C. albicans* cells did not move, but bacteria moved passively with flowing water due to their much smaller size. At around 40 min after adding SWE, the vacuole of one *C. albicans* cell collapsed, as shown in Fig. 5E-F. Yeast vacuole collapse, or fragmentation and disintegration, is a key morphological change associated with yeast cell death, often induced by antimicrobial treatment, specifically antifungal agents that target the cell membrane (Ogita et al. 2007). In a further effort to get a grasp of cell lysis, *C. albicans* cells in a growing phase were concentrated and resuspended in 10 μL fresh LB broth, 5 μL of which was loaded onto a slide. Then, 3 μL of 200 mg/mL SWE was added to the cell suspension, covered with the coverslip before being quickly loaded onto the microscope. A view of cells in hyphal form was chosen, as shown in Fig. 5G-H. Between 7 min to 10 min after adding SWE, one cell disappeared, indicating that cell lysis occurred under the action of SWE.

### SOS response regulon, sporulation regulon and ClpP govern *B. subtilis* survival against SWE treatment

Fig. 5 clearly shows that SWE kills bacteria and fungus only partially through damaging cell envelopes, assumably disrupting cell membranes, because the number of red cells does not count for the dramatic decrease in CFU. A large proportion of the cells were presumably killed through other mechanisms which could be DNA damage, DNA replication inhibition, protein synthesis inhibition, etc. Damages by such mechanisms usually induce various responses such as SOS response, sporulation activation, TA modules activation and so on. To explore the effect of such responses on the survival of *B. subtilis* upon SWE treatment, the MICs of the corresponding stress response regulator deletion mutants Δ*lexA,* Δ*abrB* and Δ*clpP* were tested and the results ware shown in Fig. 6. All regulator deletion mutants were more vulnerable to SWE treatment, especially the *clpP* deletion mutant.

**Fig. 6.**
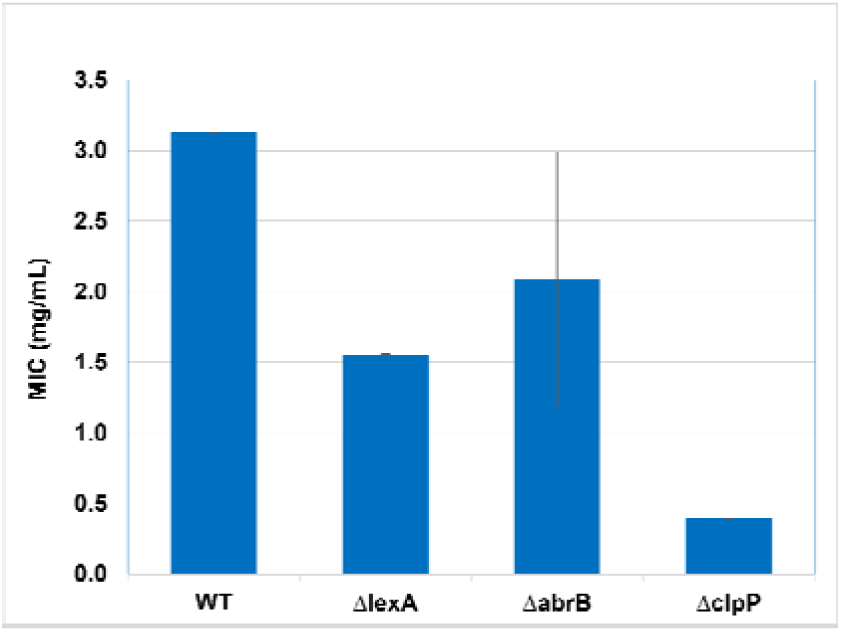
MICs of SWE to *B. subtilis* mutants Δ*lexA,* Δ*abrB* and Δ*clpP*. Error bar: standard deviation (SD, n=3)

### Co-culturing mushrooms with bacteria promoted mushroom growth

Commercial edible mushrooms have antimicrobial activities (Fig. 1). In industry they are usually grown on well controlled substrates but in nature they grow in an environment with various competitors. Here we would like to check if co-culturing commercial mushrooms with bacteria would induce stronger mushroom antimicrobial activities. Some bacteria in this study were pathogenic species and not commonly seen in nature. It was unclear if the bacteria would hinder the development of mushroom mycelium. Therefore, it was safer to introduce the bacteria into a substrate with well-established mycelium. Commercial growth kits were used for this purpose.

As shown in Fig. 1, shiitake had a strong antimicrobial effect on the tested bacteria, any subtle changes resulting from co-culturing may thus be difficult to detect. Commercial button mushroom cultivation needs a nutrient-rich compost made of pasteurized manure, making it less attractive to use button mushrooms for co-cultivation since they are usually grown with bacteria already. Oyster and lion’s mane mushrooms exhibited weak antibacterial properties (Fig. 1). This makes them good candidates for the co-culture experiment, as even minor improvements will be easily detectable. Meanwhile, they are known to be fast-growing, and their growth kits were easily commercially available. Therefore, they were chosen for co-culture experimentation.

Lion’s mane mushrooms grew healthy in all settings, as seen in the supplementary material Fig. S3A, and the weight of harvest varied slightly without a specific pattern (Fig. 7C), suggesting that the bacteria did not compromise the production of mushrooms. Instead, the mushrooms co-cultured with bacteria had hyphal knots and pins appeared earlier than the control, indicating that the bacteria promoted the development of the fruiting body of lion’s mane mushrooms (Fig. 7A).

**Fig. 7.**
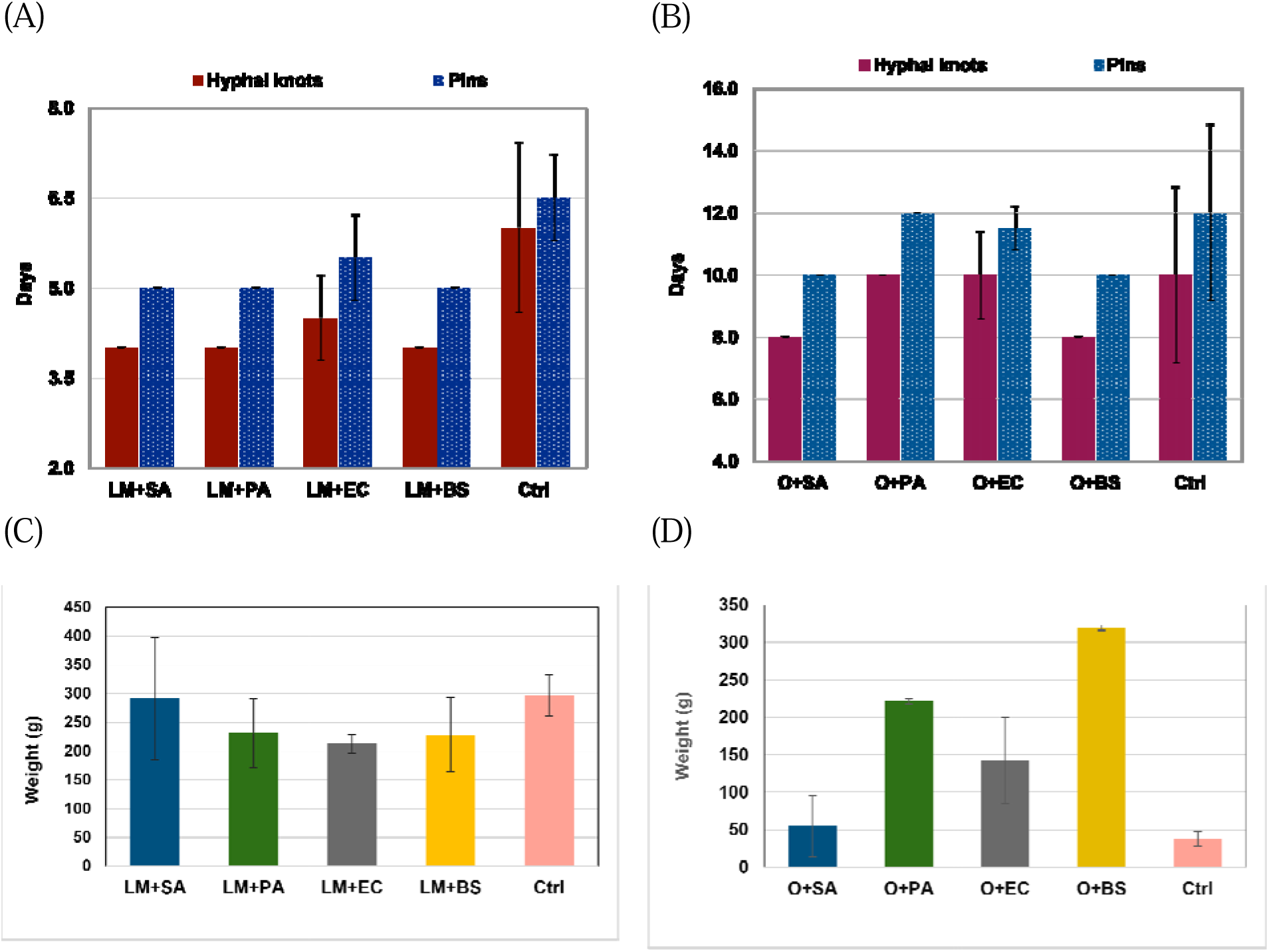
Co-culturing lion’s mane (LM) and oyster mushrooms (O) with *S. aureus* (SA)*, P. aeruginosa* (PA)*, E. coli* (EC) and *B. subtilis* (BS). “Ctrl” means water instead of bacteria was injected into mushroom substrate for co-culture. (A) The days it took for the hyphal knots and pins of lion’s mane mushroom to appear after the substrate was cut; (B) The days it took for the hyphal knots and pins of oyster mushroom to appear after the substrate was cut; (C) Weight of the harvested lion’s mane mushrooms. (D) Weight of the harvested oyster mushrooms.

In comparison to lion’s mane, the growth of oyster mushrooms was much slower and varied more significantly with different settings. The controls, where only water was injected into the mushroom substrate, performed poorly in comparison to the co-cultured ones in all tested indexes. Their hyphal knots and pins showed up later than those co-cultured with *S. aureus*, *E. coli* and *B. subtilis* and no earlier than the one co-cultured with *P. aeruginosa*. The growth of their fruiting body was very slow, with small caps and lengthy stems (Fig. S1B). The yield for the control in oyster mushrooms was the lowest among all settings. The cultivation condition was not optimal for oyster mushrooms in the study. However, the condition was the same for all oyster mushroom-bacteria co-cultures. The co-cultures overall outperformed the culture of oyster mushroom alone, indicating that these bacteria promoted the growth of oyster mushrooms.

Among the co-cultured oyster mushrooms, the growth differed significantly with different bacteria (Figs. S1B, 7B and 7D). The oyster mushrooms co-cultured with *B. subtilis* started to pin earliest, grew the healthiest, and gave the highest yield. Oyster mushrooms co-cultured with *P. aeruginosa* had the second-best yield although they pinned slightly later than other co-cultures. The oyster mushrooms co-cultured with *S. aureus* started growing early but the growth of fruiting body slowed down later. The appearance of their fruiting body was not as healthy as those co-cultured with *B. subtilis* and *P. aeruginosa,* and the yield was lower as well.

### Mushroom-bacteria coculture modestly promoted mushrooms’ antimicrobial effects

The harvests of the co-cultured mushrooms were dried at low temperature, grounded to powder from which the ethanol and water extracts were prepared and subjected to the diffusion disk susceptibility test as described before. No antimicrobial effect was observed from any water extracts. The antimicrobial effects of the ethanol extracts are shown in Fig. 8. For lion’s mane, only the co-culture with *S. aureus* improved the antimicrobial property of the ethanol extract, and such improvement was observed only in the killing against *E. coli* and *B. subtilis*, not to *S. aureus and P. aeruginosa* which are more clinically important. The co-culture with *P. aeruginosa, E. coli* and *B. subtilis* generated no visible effect (Fig. 8A).

**Fig. 8.**
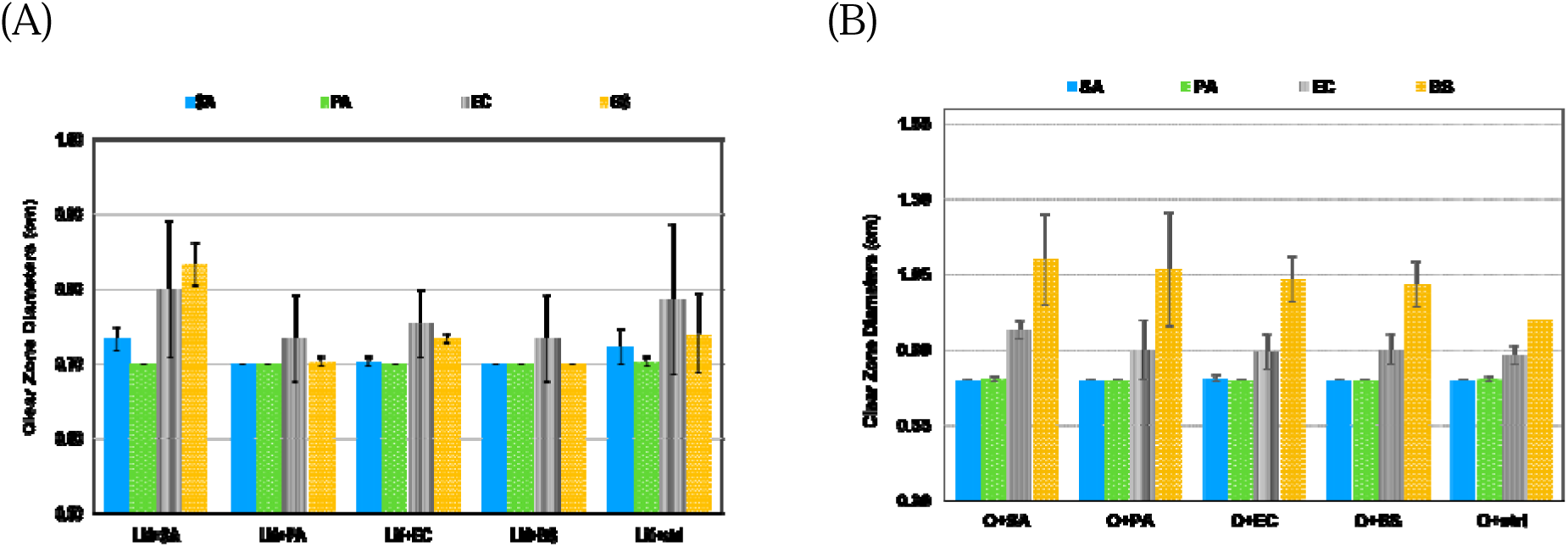
Diffusion disk susceptibility test of ethanol extracts of co-cultured mushrooms against *S. aureus, P. aeruginosa, E. coli and B. subtilis*. (A) Lion’s mane-bacteria co-culture; (B) Oyster mushroom-bacteria co-culture. Error bar: standard deviation (SD; n=3)

For oyster mushrooms, all co-cultures exhibited stronger antimicrobial effects on *E. coli* and *B. subtilis,* though more significantly on *B. subtilis* as shown in Fig. 7B. Again, co-culturing with *S. aureus* showed the greatest improvement in the antimicrobial effect. All co-cultures displayed no improvement in the killing to *S. aureus* and *P. aeruginosa* which is similar to the lion’s mane co-culture results.

## Discussion

### Characterization of SWE antimicrobial activities

Among the four tested mushrooms’ extracts, SWE exhibited the strongest antimicrobial effect against all tested bacteria and antifungal activity against *C. albicans*, but the shiitake ethanol extract had minimal effect against *B. subtilis* (Fig. 1). The broad-spectrum antimicrobial property of shiitake mushroom has been reported before. It varied significantly depending on the extraction methods and solvents, and the antimicrobial potency was attributed mainly to organic solvent extraction such as ethyl acetate or chloroform extractions (Sutthisa et al. 2025; Chasse 2026). This study followed the dual extraction method for functional food but used a cell disruptor to release cell content directly to solvents instead of the traditional soaking method. In contrast to the traditional hot water extraction, here during the process the sample container was kept in an ice bath, and the water extract was lyophilized to protect the bioactive compounds as much as possible. The result in Fig. 4A clearly showed that the antimicrobials in SWE are heat labile, losing activity before 70°C. The extraction method using hot water could cause the loss of antimicrobial effects, and so could cooking procedures. Results in Fig. 4B showed that the antimicrobials in SWE favor acidic conditions, although activity stops at very low pH conditions. The antimicrobials would likely be able to survive stomach conditions.

SWE, as a crude extract, is a cocktail of various antimicrobial compounds which are either anti-bacterial, anti-fungal or both. The MIC and MBC of SWE against each bacterium and the corresponding biofilm served as an indicator of the overall potency. By using the crystal violet staining method, the biofilm assay here focused more on measuring the mass of the biofilm. In this study, SWE was more effective against Gram-positive bacteria, no matter in inhibition or eradication, to the cells or to the biofilm. The biofilm of *B. subtilis* is an exception because it forms a thick, dense film on top of the medium surface, unlike other tested bacteria which have biofilms form on the wall of the plate wells. It is noteworthy that SWE inhibited and killed *P. aeruginosa* efficiently as shown in Figs. 2-3. *P. aeruginosa* PAO1 is known to be intrinsically resistant to almost all major antibiotic classes. Its outer membrane is tight and dense to block large, hydrophobic molecules such as macrolides, lincosamides and glycopeptides from entering the cells. It also has efflux pumps to pump out smaller antibiotics, native β-lactamase enzymes like AmpC to hydrolyze penicillins and forms highly resilient biofilms (Hu et al. 2025). Polymyxins, as a last-line treatment to *P. aeruginosa* infection, physically disrupts its outer membrane. SWE in this research probably kills *P. aeruginosa* through a similar strategy-damaging the envelope of microbial cells as indicated by Fig. 5. It is also encouraging to see that SWE inhibited the biofilm formation of *P. aeruginosa* which is the leading cause of persistent infections on medical devices and the stubborn chronic infections like cystic fibrosis. It failed in eradicating the pre-formed biofilm of *P. aeruginosa*, but was efficient in eradicating the biofilm of *S. aureus* and *E. coli,* which frequently form on host tissues and indwelling devices like catheters, prosthetic joints and heart valves.

SWE also exhibited strong antifungal effects (Fig. 5E-H). It could lyse *C. albicans* cells completely or make the vacuole collapse. When the vacuole collapses, *C. albicans* loses its ability to recycle nutrients, degrade toxic cytoplasmic waste and properly regulate intracellular calcium, ultimately leading to cell death (Palmer et al. 2005; Palmer 2002). The high potency of SWE against *C. albicans* and its food origin make SWE a great candidate for treating oral candidiasis. In addition, the antifungal activity of SWE makes it highly promising for agriculture, where fungal pathogens cause significant crop losses.

### Some antimicrobial compounds in SWE are likely proteins and glycans

Bioactive compounds from shiitake mycelium and fruiting body can be found in Scientific, Natural Product, and food databases. Antimicrobial compounds from shiitake were included in but not limited to Supplementary Table S2. They were reportedly enriched in organic solvent extracts, such as the well-known lenthionine and lentinamycin (Morita et al. 1967; Hirasawa et al. 1999; Shimada et al. 2024; Chasse 2026). Isolating and characterizing each of them would be technically challenging. In this study the significant antimicrobial activities of shiitake were from the water extract. The possible water-soluble antimicrobial compounds were analyzed here and some experiments were carried out to explore the contributors.

Dianhydromannitol (Supplementary Table S2) is soluble in both ethanol and water, and was reported to be one of the major components in shiitake ethanol extract exhibiting antibacterial effect (Erdoğan Eliuz 2022). In this study, ethanol extraction was carried out first and the residue was subjected to water extraction. Dianhydromannitol was thus expected to be mainly in the ethanol extract of shiitake. However, the shiitake ethanol extract did not show much antibacterial effect (Fig. 1). If it was present in SWE in our research, it would not be the main contributor of antimicrobial activity since it is heat stable but SWE is heat labile (Fig. 4A).

Lentin is a 27.5KDa water-soluble antifungal and antiviral protein in the fruiting body of shiitake (Baral B. 2025; Ngai et al. 2003; Minutti et al. 2016). In addition, an unnamed protein of 87.2 KDa purified from the aqueous extract of shiitake exhibited antimicrobial effects against *E. coli and S. aureus* (Minutti et al. 2016). Here the protease digestion reduced the clear zone of diffusion disk on *E. coli* lawn (Fig. 4C), indicating that some of the antimicrobials were proteins. Lentin was specifically tested against fungi and virus but not tested for bacteria. The unknown protein of 87.2 KDa was tested against bacteria but not fungi. SWE in this study was effective against both bacteria and fungi, probably with contributions from these two proteins.

Lentinan is the major water-soluble component from shiitake and clinically approved in Japan and China as an immunostimulant for the treatment of various cancers. It was also reported to have an antimicrobial effect indirectly from its immunity-boosting property (Liu et al. 2019). The dose-dependent effect of β-glucanase digestion on the dwindling inhibition zone of diffusion disk in Fig.4 D strongly indicated that lentinan can kill bacteria directly. To verify this, it would be ideal to use pure lentinan. In practice, the bioactivity of lentinan manufactured for agriculture applications is sensitive to temperature and pH, being stable when the environment is neutral to slightly acidic (a pH between 5.5 and 7.0) or under 70°C (Green Agri Bio, 2024), which is consistent with the experimental results shown in Fig. 4A-B. However, its purity is low. The commercial lentinan manufactured for research or clinical purposes has a high purity. However, it is usually prepared through hot water and alkaline extraction, alcohol precipitation, and multi-stage purification with chromatography, etc. In their natural state, β-glucans like lentinan exist as rigid, self-stabilizing triple helices. Harsh purification processes—like the use of strong alkaline solutions or DMSO—break the hydrogen bonds holding the helices together, causing them to denature into single-strand random coils. The renaturation results in a mixture of linear, circular and branched species of triple helix, causing the loss of or decreased bioactivities (Caseiro et al. 2022). Therefore, the antimicrobial activity of commercial lentinan with a high purity may have been greatly reduced. Nevertheless, lentinan with a purity above 99% was purchased from GLPBio (Good Laboratory Practice Bioscience), made into final concentration of 30 mg/mL at pH 5, 6, 7.5, 8 and 9, respectively, and spotted on paper diffusion disk to test against all four bacteria and *C. albicans*, but no antimicrobial effect was observed (data not shown). A better purification method is thus indispensable for the verification of the direct antimicrobial effect of lentinan.

### Killing mechanisms of antimicrobials in SWE

Among the tested microbes, SWE was the most efficient against *B. subtilis* and *C. albicans.* Only a fraction of their dead cells had the envelope damaged by SWE, indicating the existence of other killing mechanisms (Fig. 5A-D). Lentinan was reported to inhibit the activity of DNA topoisomerase I *in vitro* in a dose-dependent manner (Yehia 2022). Edodin and ledodin, which inhibit protein synthesis, are ribosome-inactivating proteins isolated from shiitake, but they only work against mammalian ribosomes (Citores et al. 2023 and 2024). These large molecules would need to get into the microbial cells first before they can target the DNA-interacting enzymes or protein synthesizing ribosomes. Considering this, the small molecules such as benzoic acid derivatives, quinolones, oxalic, succinic, quinic acids, adenine, inosine and uridine etc. may be responsible for the killings through mechanisms other than envelope damaging (Papetti et al. 2018; Baral 2025). However, these small molecules are relatively heat stable which does not align well with the results shown in Fig. 4A.

Apparently, without isolating and purifying the specific antimicrobials from SWE, it would be hard to clearly elucidate the killing mechanisms from the origin. However, the sharp drop in MICs to Δ*lexA,* and Δ*clpP* mutants implies that SWE damages DNA and inhibits protein synthesis as well (Fig. 6). The possible mechanism reflected by the decreased MIC to Δ*abrB mutant* would be more specific to *B. subtilis*.

As mentioned before, the genome sequencing of the *B. subtilis* mutant resistant to SWE generated a list of mutated genes in the regulons of sporulation, quorum sensing, PBSX prophage, SPβ prophage, DNA repair and TA modules (Supplementary Table S1). These regulons are usually repressed under healthy growth conditions but induced upon stresses to either help the cells endure the stresses by repairing DNA, remove proteins of malfunction, form spores, kill other invading microbes or initiate cannibalism for the survival of the population.

Deleting the regulators in these regulons would interfere with cell survival against SWE treatment, providing some clues of the SWE killing mechanisms. For instance, AbrB protein, the global transition state regulator under the master regulator Spo0A, represses the expression of autolysin *cwlH —* a peptidoglycan hydrolase damaging the cell wall, and the large *srfA* and *ppsABCDE* regulons which synthesize surfactin and plipastatin, respectively, both of which fight competing microbes (Vahidinasab et al. 2020). Upon SWE treatment, Spo0A may reverse the repression by inhibiting the *abrB* expression and indirectly inducing protease ClpP to cleave AbrB, subsequently initiating hydrolysis of cell wall and unnecessarily expressing the *srfA* and *ppsABCDE* regulons. Deleting *abrB* would permanently turn on the expression of autolysin cwlH, damaging the cell wall and allowing more antimicrobials to enter cells easily. Meanwhile, constantly expressing the large *srfA* and *ppsABCDE* regulon would exhaust the cells, making them more vulnerable. Indeed, the mutant Δ*abrB* was more susceptible to SWE treatment (Fig. 6). LexA is the master repressor of SOS response. DNA damage activates RecA, initiating the self-cleavage of LexA to induce SOS response. The ClpXP protease complex then rapidly degrades these cleaved fragments of LexA to prevent the repressor from reassembling and allows the SOS-regulated genes to remain active until DNA damages are repaired. On the other hand, in *B. subtilis,* SOS response triggers the induction of prophages SPβ and PBSX, leading to the production of holins and endolysins that make pores in the cell membrane and degrade the peptidoglycan cell wall, causing cell lysis (Longchamp et al. 1994; Catalão 2013; Toyofuku 2017). At first glimpse, deleting *lexA* would keep SOS response constantly on, making the cells more resistant to DNA damaging agents. However, SOS response is a complicated global regulation network. Keeping it constantly on may put a huge burden on the cells. In addition, it would keep activating the prophages to produce holins and endolysins to kill the cells. This would explain why Δ*lexA* mutant was two times more sensitive than the wild type (Fig. 6). The effect of deleting *clpP* would be more complicated. ClpP (Caseinolytic protease P) works alongside chaperone proteins ClpC or ClpX to clear misfolded or damaged proteins and destroy specific regulatory proteins to create the quality-control system and regulate the responses to various stresses of the bacteria, such as sporulation, SOS response, competence, biofilm formation, and oxidative stress etc. (Mo et al. 2010; Msadek et al. 1998; Capestany et al. 2008; Roy et al. 2019). In addition, in *B. subtilis,* ClpP is the primary molecular trigger that activates Type II TA modules, a two-component system consisting of a highly unstable antitoxin and a stable toxin. The antitoxin in general tightly binds to the toxin to make it harmless. During stress, ClpP shreds the antitoxin to release the toxin which slows down or completely stops essential cell functions such as cell division, cell wall synthesis, DNA replication, and protein synthesis. Deleting *clpP* will allow for the build-up of damaged proteins and compromise the capability of bacterial cells to cope with stresses, so assumably it will make cells more susceptible to SWE treatment. On the other hand, deleting *clpP* would potentially reduce the accumulation of the toxins from TA modules and the activation of holins, endolysins, prophages, etc., increasing the tolerance to SWE treatment. Deleting *clpP* drastically decreased the MIC of SWE to *B. subtilis* by 8-fold (Fig. 6), indicating overall ClpP plays a significant role in *B. subtilis’* survival to SWE treatment.

### Co-culturing with bacteria promoted mushroom growth

This research showed that co-culturing with bacteria boosted the growth of both lion’s mane and oyster mushrooms. Lion’s mane is said to be fragile and slow growing, meaning its substrate must be strictly sterilized to eliminate competitors. Oyster mushrooms grow fast and are versatile with their substrate which could be either pasteurized or sterilized when mixed with high nutrient ingredients. It turned out in this study that lion’s mane in all mushroom-bacteria co-culture conditions grew healthy and was resilient to the fluctuation of growth conditions. The performance of oyster mushrooms in the same growth tent, in contrast, varied depending on the mushroom-bacteria pair and maybe other unknown factors. All mushroom-bacteria co-culture settings facilitated the growth of mushrooms, either making the hyphal knot form earlier or pins develop faster. In the past, the research on the association between bacteria and mushroom growth focused on the cultivation of white button mushrooms which need fermented compost and a casing layer for cultivation. However, research about bacteria and mushrooms growing together has expanded to various other mushrooms recently (Shamugam et al. 2023). It has been established that specific bacteria help mushroom growth in a few ways. First, bacteria help with nutrient supply. Mushroom mycelia produce bacteriolytic enzymes to process bacteria as a source of nutrients. Some bacteria process substances in substrates, such as cellulose, to turn them into consumable nutrients for mushrooms. Secondly, some bacteria can remove the inhibitory compounds produced by the mycelium. For instance, *Pseudomonas putida* inhibited the ethylene biosynthetic pathway of *Morchella sextelata* by producing 1-aminocyclopropane-1-carboxylic acid (ACC) deaminase (AcdS), and in doing so relieved the mycelium growth inhibition by ethylene (Zou et al. 2025). *Serratia rubidaea*, *Klebsiella pneumoniae* and *Bacillus cereus* presented within the casing layer of *A. bisporus* cultivation were also reported to lower the ethylene level by producing AcdS to cleave ACC, the precursor of ethylene (Chen et al. 2013). The *pseudomonad* populations in the casing were reported to consume the inhibitory C8 compounds produced by the mycelium and substrate of *A. bisporus*, promoting primordium formation (mushroom pinning) (Noble et al. 2009). Thirdly, bacteria such as *Pseudomonas* and *Bacillus* produce low-molecular-weight volatile organic compounds (VOCs) that diffuse through the substrate, signaling the fungus to initiate pinning (primordia formation) (Orban et al. 2023). Lastly, some bacteria indirectly help mushroom growth by fighting against other bacteria pathogenic to the mushrooms, such as *Pseudomonas tolaasii* which can cause bacterial blotch (Hermenau et al. 2020).

Most research on the relationship between mushrooms and bacteria focused on the natural species in soil. In this study, among the four chosen bacteria, *B. subtilis* and *P. aeruginosa* are natural soil bacteria, although *P. aeruginosa* is also a notorious opportunistic pathogen causing significant clinical concerns. Both *S. aureus* and *E. coli* are not soil bacteria. *S. aureus* is an opportunistic pathogen that colonizes the skin and upper respiratory tract of human beings or animals, and *E. coli* primarily habitats in the lower intestines of warm-blooded mammals. Surprisingly, *S. aureus* and *E. coli* also promoted the hyphal formation and pinning of both mushrooms. This indicates that such bacterial benefits to mushroom growth might be a very common phenomenon.

The growth of the oyster mushroom controls appeared to be inhibited by unidentified variables. They pinned late, and became ill soon after pinning, producing yellowish and bent fruiting bodies with a rough surface and low yield. The inhibitory effect was alleviated by bacterial co-culture. All bacteria promoted oyster mushroom development and yield, with *B. subtilis* as the best performer, followed by *P. aeruginosa* (Fig. 7). In commercial mushroom cultivation, *B. subtilis* is generally considered a beneficial bacterium, helping create a healthy microbiome for oyster mushrooms. It can inhibit pathogenic fungi such as *Trichoderma pleuroti* and *Trichoderma pleuroticola* by 54–69%, such that it can be used for controlling green mold disease in oyster mushroom production (Potocnik et al. 2019). Also, it can induce the expression of laccase enzymes that break down lignin, improving the growth and health of the *P. ostreatus* mycelium (Velazquez-Cedeno et al. 2008). The correlation between *Pseudomonas* bacteria and mushroom growth is complicated, depending on the specific bacterial strain and the mushroom species involved. *P. aeruginosa* inhibited the hyphal growth of button mushrooms upon direct contact with the mycelium; furthermore, it severely reduced mushroom yield when inoculated into the casing soil (Anwar et al. 2015). In contrast, inoculating cultures with *Pseudomonas sp*. P7014 significantly increased the growth and yield of mushrooms like *Pleurotus eryngii* (King Oyster) (Kim et al. 2007). The relationship between *P. aeruginosa* and oyster mushrooms was not well-documented in scientific literature. In a sharp contrast to the severe reduction in the yield of button mushroom by the inoculation of *P. aeruginosa* into the casing soil, in this study the co-culture of oyster mushroom with *P. aeruginosa* generated the second-best yield among all settings (Fig. 7D) and enormous fruiting bodies during the second flush (Supplementary Fig. S4).

In this study the substrates for mushroom cultivation were from commercial mushroom grow kit, with mycelium pre-established. In future, it would be interesting to make a thorough examination on the relationship between these bacteria, especially *S. aureus* and *E. coli,* and lion’s mane or oyster mushrooms, based on well-defined, fresh-made sterile substrates, and starting from the interaction between bacteria and mushroom mycelium. Bacterial and fungal metabolites and VOCs shall be examined to explore the mechanisms accounting for the mushroom growth promotion.

### Co-culturing with bacteria enhanced mushroom antimicrobial activities

This study found that mushroom-bacteria co-culture did improve mushroom antimicrobial activity, but in a broad-spectrum manner, not necessarily against the paired bacteria (Fig. 8). In a natural environment, fungi and bacteria have interactions ranging from antagonism to mutualism, creating varying effects on nutrition, growth, stress resistance and pathogenicity to hosts (Deveau et al. 2018; Zhou et al. 2022). When a fungus detects a foreign bacterium, it launches chemical defenses, including many secondary metabolites such as penicillin through stress responses to fight against a large range of bacteria rather than viewing the stimulating bacteria as its primary threat. Meanwhile, the specific bacterium that triggered the fungal stress responses may have already developed resistance to antimicrobials from that fungus. All these are consistent with the findings in this study (Fig. 8). *B. subtilis* increased oyster mushroom’s killing against itself.

Among all the co-culture settings, *S. aureus* increased lion’s mane’s antimicrobial effect on *B. subtilis* more than itself and increased oyster mushroom’s antimicrobial potency on *E. coli* and *B. subtilis* instead of itself. *P. aeruginosa,* which inhabits primarily soil, increased oyster mushroom’s killing to *E. coli* and *B. subtilis*, but *P. aeruginosa* itself was resistant to oyster mushroom extracts of all settings in this study. Although no specific co-culture pair from this study has been scientifically reported before, some research showcased a similar effect of triggering a heightened response in fungi against bacteria it was not co-cultured with. For instance, co-culturing fungus *Aspergillus sp.* CO2 with bacterium *Bacillus sp.* COBZ21 increased the antimicrobial activity of the fungus against *E. coli* ATCC 25922, *S. aureus* NRRLB-767 and *C. albicans* ATCC 10231 (Hamed et al. 2024). Also, the co-cultivation of some soil fungi with *S. aureus, B. subtilis* and *E. coli* induced copious production of some broad-spectrum antibiotics and the generation of some novel antibacterial secondary metabolites (Kenneth et al. 2017).

Notably, among the four bacteria, *S. aureus* performed the best in enhancing the antimicrobial properties of both mushrooms. This might be due to its “foreigner” status to natural environments since it usually lives on the skin and respiratory systems of human beings and animals, although it also prevails with seasonal fluctuation in some natural environments with frequent human activities (Thapaliya et al. 2017). *S. aureus* was the only bacterium that increased lion’s mane antimicrobial activity, although the increase was much slighter in comparison to its contribution to oyster mushroom (Fig. 7). For the oyster mushroom, all bacteria improved the mushroom antimicrobial ability to *E. coli* and *B. subtilis,* but *S. aureus* contributed the most.

This study focused on the edible part of the mushroom—fruiting bodies. To further study the mechanism of the production of mushroom antimicrobials upon bacterial stimulation, it would be more interesting to focus on the mycelium since it interacts with bacteria and the substrate directly.

## Conclusion

Among the four tested edible mushrooms from the local market, shiitake exhibited the strongest antimicrobial effect, followed by button, lion’s mane and oyster mushroom.

Antimicrobials from shiitake kill bacteria and fungi and inhibit bacterial biofilms. They kill microbes through damaging cell envelope, lysing cells, and probably damaging DNA and inhibiting protein synthesis as well. Some of the antimicrobials are possibly proteins and polysaccharides. Lentinan from shiitake could be able to kill microbes directly, rather than indirectly through modulating host immune systems. Obtaining lentinan with a high purity without loss of bioactivity is the key to verifying its direct microbial killing ability. Mushroom-bacteria co-cultures overall were beneficial to mushroom growth and enhanced mushroom antimicrobial property in a broad-spectrum manner, indicating its potential in both agriculture and medicinal application.

## Acknowledgements

We would like to thank the Massachusetts Science and Engineering Fair (MSEF) as this project was initiated as an entry to the fair.

## Author contributions

Eunice Wang conceptualized the study, performed the experiments, analyzed the data and wrote the manuscript. Nicole Cavanaugh and Yinghao He aided in carrying out experiments. Yunrong Chai provided supervision, guidance, and review of the manuscript.

## References

1. Alves, M., Ferreira, I., Dias, J., Teixeira, V., Martins, A., & Pintado, M. (2012). A review on antimicrobial activity of mushroom (basidiomycetes) extracts and isolated compounds. Planta Medica, 78(16), 1707–1718. 10.1055/s-0032-1315370

2. Al Qutaibi, M. & Kagne, S. R. (2024). Unearthing nature’s pharmacy: Exploring the antimicrobial potency of mushrooms. Journal of Food Processing and Preservation, 2024(1). 10.1155/2024/8331974

3. Anke T., Oberwinkler, F., Steglich, W., & Schramm, G. (1977). The strobilurins - new antifungal antibiotics from the basidiomycete *Strobilurus tenacellus*. The Journal of Antibiotics, 30(10), 806–810. 10.7164/antibiotics.30.806

4. Baral, B. (2025). Holistic evaluation of shiitake mushrooms (*Lentinula edodes*): Unraveling its medicinal and therapeutic potentials. Chemistry & Biodiversity, 22(12), e01244. 10.1002/cbdv.202501244

5. Capestany, C. A., Tribble, G. D., Maeda, K., Demuth, D. R., & Lamont, R. J. (2008). Role of the CLP system in stress tolerance, biofilm formation, and intracellular invasion in *Porphyromonas gingivalis*. Journal of Bacteriology, 190(4), 1436–1446. 10.1128/jb.01632-07

6. Caseiro, C., Dias, J. N. R., de Andrade Fontes, C. M. G., & Bule, P. (2022). From cancer therapy to wine making: The molecular structure and applications of β-Glucans and β-1, 3-Glucanases. International Journal of Molecular Sciences, 23(6), 3156. 10.3390/ijms23063156

7. Catalão, M. J., Gil, F., Moniz-Pereira, J., São-José, C., & Pimentel, M. (2013). Diversity in bacterial lysis systems: Bacteriophages show the way. FEMS Microbiology Reviews, 37(4), 554–571. 10.1111/1574-6976.12006

8. Chasse, J. (2026). Chromatographic Profiling and Antibacterial Activity of Solvent-Extracted Shiitake Mushroom Compounds. Chromatographyonline.Com. https://www.chromatographyonline.com/view/chromatographic-profiling-and-antibacterial-activity-of-solvent-extracted-shiitake-mushroom-compounds

9. Chen, S., Qiu, C., Huang, T., Zhou, W., Qi, Y., Gao, Y., Shen, J., & Qiu, L. (2013). Effect of 1-aminocyclopropane-1-carboxylic acid deaminase producing bacteria on the hyphal growth and primordium initiation of *Agaricus bisporus*. Fungal Ecology, 6(1), 110–118. 10.1016/j.funeco.2012.08.003

10. Citores, L., Ragucci, S., Gay, C. C., Russo, R., Chambery, A., Di Maro, A., Iglesias, R., & Ferreras, J. M. (2024). Edodin: A new type of toxin from shiitake mushroom (*Lentinula edodes*) that inactivates mammalian ribosomes. Toxins, 16(4), 185. 10.3390/toxins16040185

11. Citores, L., Ragucci, S., Russo, R., Gay, C. C., Chambery, A., Di Maro, A., Iglesias, R., & Ferreras, J. M. (2023). Structural and functional characterization of the cytotoxic protein ledodin, an atypical ribosome-inactivating protein from shiitake mushroom (*Lentinula edodes)*. Protein Science : A Publication of the Protein Society, 32(4), e4621. 10.1002/pro.4621

12. Deveau, A., Bonito, G., Uehling, J., Paoletti, M., Becker, M., Bindschedler, S., Hacquard, S., Hervé, V., Labbé, J., Lastovetsky, O. A., Mieszkin, S., Millet, L. J., Vajna, B., Junier, P., Bonfante, P., Krom, B. P., Olsson, S., van Elsas, J. D., & Wick, L. Y. (2018). Bacterial–fungal interactions: Ecology, mechanisms and challenges. FEMS Microbiology Reviews, 42(3), 335–352. 10.1093/femsre/fuy008

13. Erdoğan Eliuz, E. A. (2022). Antibacterial activity and antibacterial mechanism of ethanol extracts of *Lentinula edodes* (shiitake) and *Agaricus bisporus* (button mushroom). International Journal of Environmental Health Research, 1–14. 10.1080/09603123.2021.1919292

14. Green Agri Bio. (2024, December 27). The importance of high-quality lentinan in agriculture. https://www.greenagribio.com/news/the-importance-of-high-quality-lentinan-in-agriculture.html

15. Hamed, A. A., Ghareeb, M. A., Kelany, A. K., Abdelraof, M., Kabary, H. A., Soliman, N. R., & Elawady, M. E. (2024). Induction of antimicrobial, antioxidant metabolites production by co-cultivation of two red-sea-sponge-associated Aspergillus sp. CO2 and Bacillus sp. COBZ21. BMC Biotechnology, 24(1), 3. 10.1186/s12896-024-00830-z

16. Hayashi, K., Morooka, N., Yamamoto, Y., Fujita, K., Isono, K., Choi, S., Ohtsubo, E., Baba, T., Wanner, B. L., Mori, H., & Horiuchi, T. (2006). Highly accurate genome sequences of escherichia coli K-12 strains MG1655 and W3110. Molecular Systems Biology, 2, 2006.0007. 10.1038/msb4100049

17. Hermenau, R., Kugel, S., Komor, A. J., & Hertweck, C. (2020). Helper bacteria halt and disarm mushroom pathogens by linearizing structurally diverse cyclolipopeptides. Proceedings of the National Academy of Sciences, 117(38), 23802–23806. 10.1073/pnas.2006109117

18. Hirasawa, M., Shouji, N., Neta, T., Fukushima, K., & Takada, K. (1999). Three kinds of antibacterial substances from *Lentinus Edodes* (Berk.) Sing. (Shiitake, an edible mushroom). International Journal of Antimicrobial Agents, 11(2), 151–157. 10.1016/s0924-8579(98)00084-3

19. Hu, M., & Chua, S. L. (2025). Antibiotic-resistant *Pseudomonas aeruginosa*: Current challenges and emerging alternative therapies. Microorganisms, 13(4), 913. 10.3390/microorganisms13040913

20. Jones, T., Federspiel, N. A., Chibana, H., Dungan, J., Kalman, S., Magee, B. B., Newport, G., Thorstenson, Y. R., Agabian, N., Magee, P. T., Davis, R. W., & Scherer, S. (2004). The diploid genome sequence of candida albicans. Proceedings of the National Academy of Sciences, 101(19), 7329–7334. 10.1073/pnas.0401648101

21. Kenneth GN, Nwoye VO, Okoye FBC, Proksch P. (2017). Staphylococcus aureus, Bacillus subtilis and Escherichia coli induced copious production of antibiotics in an overnight co-culture with three soil fungi. Journal of current Biomedical Research. 1(1):65–72

22. Kim, M.K., Math, R.K., Cho, K.M., Shin, K.J., Kim, J.O., Ryu, J.S., Lee, Y.H., Yun, H.D. (2007). Effect of Pseudomonas sp. P7014 on the growth of edible mushroom Pleurotus eryngii in bottle culture for commercial production. Bioresour Technol. 99(8):3306-8.

23. Liang, L., Su, Q., Ma, Y., Zhao, S., Zhang, H., & Gao, X. (2024). Research progress on the polysaccharide extraction and antibacterial activity. Annals of Microbiology, 74(1). 10.1186/s13213-024-01762-x

24. Liu, Y., Zhao, J., Zhao, Y., Zong, S., Tian, Y., Chen, S., Li, M., Liu, H., Zhang, Q., Jing, X., Sun, B., Wang, H., Sun, T., & Yang, C. (2019). Therapeutic effects of lentinan on inflammatory bowel disease and colitis-associated cancer. Journal of Cellular and Molecular Medicine, 23(2), 750–760. 10.1111/jcmm.13897

25. Longchamp, P. F., Mauël, C., & Karamata, D. (1994). Lytic enzymes associated with defective prophages of Bacillus subtilis: Sequencing and characterization of the region comprising the N-acetylmuramoyl-L-alanine amidase gene of prophage PBSX. Microbiology (Reading, England), 140 (Pt 8), 1855–1867. 10.1099/13500872-140-8-1855

26. Minutti, L., Téllez-Téllez, M., R, D., S, T.-B., FJ, F., Santos-López, G., & Diaz-Godínez, G. (2016). Antimicrobial activity of a protein obtained from fruiting body of *Lentinula edodes* against *Escherichia coli* and *Staphylococcus aureus*. Journal of Environmental Biology, 37(4), 619–623.

27. Mo, A. H., & Burkholder, W. F. (2010). YneA, an SOS-Induced inhibitor of cell division in *Bacillus subtilis*, is regulated posttranslationally and requires the transmembrane region for activity. Journal of Bacteriology, 192(12), 3159–3173. 10.1128/jb.00027-10

28. Mohammad, A., & Sabaa, A. (2015). *In vitro* and *in vivo* impact of some *Pseudomonas* spp. on the growth and yield of cultivated mushroom (*Agaricus bisporus*). The Egyptian Journal of Experimental Biology (Botany), 11(2): 163 – 167. 10.5455/esebb.20151029051037

29. Morita, K., & Kobayashi, S. (1967). Isolation, structure, and synthesis of lenthionine and its analogs. Chemical and Pharmaceutical Bulletin, 15(7), 988–993. 10.1248/cpb.15.988

30. Msadek, T., Dartois, V., Kunst, F., Herbaud, M. L., Denizot, F., & Rapoport, G. (1998). ClpP of *Bacillus subtilis*is required for competence development, motility, degradative enzyme synthesis, growth at high temperature and sporulation. Molecular Microbiology, 27(5), 899–914. 10.1046/j.1365-2958.1998.00735.x

31. Ngai, P. H. K., & Ng, T. B. (2003). Lentin, a novel and potent antifungal protein from shitake mushroom with inhibitory effects on activity of human immunodeficiency virus-1 reverse transcriptase and proliferation of leukemia cells. Life Sciences, 73(26), 3363–3374. 10.1016/j.lfs.2003.06.023

32. Noble, R., Dobrovin-Pennington, A., Hobbs, P. J., Pederby, J., & Rodger, A. (2009). Volatile C8 compounds and *Pseudomonads* influence primordium formation of *Agaricus bisporus*. Mycologia, 101(5), 583–591. 10.3852/07-194

33. Nye, T. M., Schroeder, J. W., Kearns, D. B., & Simmons, L. A. (2017). Complete genome sequence of undomesticated bacillus subtilis strain NCIB 3610. Genome Announcements, 5(20). 10.1128/genomea.00364-17

34. Ogita, A., Nagao, Y., Fujita, K.-I., & Tanaka, T. (2007). Amplification of vacuole-targeting fungicidal activity of antibacterial antibiotic polymyxin B by allicin, an allyl sulfur compound from garlic. The Journal of Antibiotics, 60(8), 511–518. 10.1038/ja.2007.65

35. Okamoto, K., Gotoh, N., & Nishino, T. (2001). *Pseudomonas aeruginosa* reveals high intrinsic resistance to penem antibiotics: Penem resistance mechanisms and their interplay. Antimicrobial Agents and Chemotherapy, 45(7), 1964–1971. 10.1128/AAC.45.7.1964-1971.2001

36. Orban, A., Jerschow, J. J., Birk, F., Suarez, C., Schnell, S., & Rühl, M. (2023). Effect of bacterial volatiles on the mycelial growth of mushrooms. Microbiological Research, 266, 127250. 10.1016/j.micres.2022.127250

37. Palmer, G. (2002). Functional analysis of the vacuole in Candida albicans. University of Leicester. Thesis. https://hdl.handle.net/2381/29046

38. Palmer, G. E., Kelly, M. N., & Sturtevant, J. E. (2005). The *Candida albicans* vacuole is required for differentiation and efficient macrophage killing. Eukaryotic Cell, 4(10), 1677–1686. 10.1128/EC.4.10.1677-1686.2005

39. Pang, Z., Raudonis, R., Glick, B. R., Lin, T.-J., & Cheng, Z. (2019). Antibiotic resistance in *Pseudomonas aeruginosa*: Mechanisms and alternative therapeutic strategies. Biotechnology Advances, 37(1), 177–192. 10.1016/j.biotechadv.2018.11.013

40. Papetti, A., Signoretto, C., Spratt, D. A., Pratten, J., Lingström, P., Zaura, E., Ofek, I., Wilson, M., Pruzzo, C., & Gazzani, G. (2018). Components in *Lentinus edodes* mushroom with anti-biofilm activity directed against bacteria involved in caries and gingivitis. Food & Function, 9(6), 3489–3499. 10.1039/c7fo01727h

41. Potocnik, I., Milijasevic-Marcic, S., Stanojevic, O., Beric, T., Stankovic, S., Kredics, L., & Hatvani, L. (2019). The activity of native *Bacillus subtilis* strains in control of green mould disease of oyster mushroom (*Pleurotus spp*.). Pesticidi I Fitomedicina, 34(2), 97–102. 10.2298/pif1902097p

42. Robertson, J., McGoverin, C., White, J. R., Vanholsbeeck, F., & Swift, S. (2021). Rapid detection of *Escherichia coli* antibiotic susceptibility using live/dead spectrometry for lytic agents. Microorganisms, 9(5), 924. 10.3390/microorganisms9050924

43. Roy, S., Zhu, Y., Ma, J., Roy, A. C., Zhang, Y., Zhong, X., Pan, Z., & Yao, H. (2019). Role of ClpX and ClpP in *Streptococcus suis* serotype 2 stress tolerance and virulence. Microbiological Research, 223*–*225, 99–109. 10.1016/j.micres.2019.04.003

44. Sassi, M., Felden, B., & Augagneur, Y. (2014). Draft genome sequence of staphylococcus aureus subsp. *aureus* strain HG003, an NCTC8325 derivative. Genome Announcements, 2(4). 10.1128/genomea.00855-14

45. Shamugam, S., & Kertesz, M. A. (2023). Bacterial interactions with the mycelium of the cultivated edible mushrooms *Agaricus bisporus* and *Pleurotus ostreatus*. Journal of Applied Microbiology, 134(1), lxac018. 10.1093/jambio/lxac018

46. Shimada, S., Yamaguchi, Y., Saki Nakajo, Mochizuki, S., Hirata, R., Tanabe, Y., Yamada, K., & Kumagai, H. (2024). Formation mechanism of an inclusion complex of lenthionine with α-cyclodextrin and enhancement of its bioavailability. Journal of Food Bioactives, 28, 68–75. 10.26599/jfb.2024.95028397

47. Stover, C. K., Pham, X. Q., Erwin, A. L., Mizoguchi, S. D, Warrener, P., Hickey, M. J., Brinkman, F. S., Hufnagle, W. O., Kowalik, D. J., Lagrou, M., Garber, R. L., Goltry, L., Tolentino, E., Westbrock-Wadman, S., Yuan, Y., Brody, L. L., Coulter, S. N., Folger, K. R., Kas, A., … Olson, M. V. (2000). Complete genome sequence of pseudomonas aeruginosa PAO1, an opportunistic pathogen. Nature, 406(6799), 959–964. 10.1038/35023079

48. Sutthisa, W., Kamlangmak, P., & Srisawad, N. (2025). Antibacterial potential and chemical composition of shiitake mushroom (*Lentinus edodes* (Berk.) Sing.) Extract against pathogenic bacteria. Scientifica, 2025(1). 10.1155/sci5/6089332

49. Thapaliya, D., Hellwig, E. J., Kadariya, J., Grenier, D., Jefferson, A. J., Dalman, M., Kennedy, K., DiPerna, M., Orihill, A., Taha, M., & Smith, T. C. (2017). Prevalence and characterization of *Staphylococcus aureus* and methicillin-resistant *Staphylococcus aureus* on public recreational beaches in northeast Ohio. GeoHealth, 1(10), 320–332. 10.1002/2017gh000106

50. Toyofuku, M., Cárcamo-Oyarce, G., Yamamoto, T., Eisenstein, F., Hsiao, C.-C., Kurosawa, M., Gademann, K., Pilhofer, M., Nomura, N., & Eberl, L. (2017). Prophage-triggered membrane vesicle formation through peptidoglycan damage in *Bacillus subtilis*. Nature Communications, 8(1), 481. 10.1038/s41467-017-00492-w

51. Vahidinasab, M., Lilge, L., Reinfurt, A., Pfannstiel, J., Henkel, M., Morabbi Heravi, K., & Hausmann, R. (2020). Construction and description of a constitutive plipastatin mono-producing *Bacillus subtilis*. Microbial Cell Factories, 19(1), 205. 10.1186/s12934-020-01468-0

52. Valverde, M. E., Hernández-Pérez, T., & Paredes-López, O. (2015). Edible mushrooms: Improving human health and promoting quality life. International Journal of Microbiology, 2015(376387), 1–14. 10.1155/2015/376387

53. Velázquez-Cedeño, M., Farnet, A. M., Mata, G., & Savoie, J.-M. (2008). Role of *Bacillus spp*. in antagonism between *Pleurotus ostreatus* and *Trichoderma harzianum* in heat-treated wheat-straw substrates. Bioresource Technology, 99(15), 6966–6973. 10.1016/j.biortech.2008.01.022

54. WHO. (2025) Antimicrobial Resistance Division (AMR). Global antibiotic resistance surveillance report 2025. In Who.int. World Health Organization. https://www.who.int/publications/i/item/9789240116337

55. Yehia, R. S. (2022). Evaluation of the biological activities of β ßglucan isolated from *Lentinula edodes*. Letters in Applied Microbiology, 75(2), 317–329. 10.1111/lam.13727

56. Zhou, Y., Wang, H., Xu, S., Liu, K., Qi, H., Wang, M., Chen, X., Berg, G., Ma, Z., Cernava, T., & Chen, Y. (2022). Bacterial-fungal interactions under agricultural settings: From physical to chemical interactions. Stress Biology, 2(1). 10.1007/s44154-022-00046-1

57. Zou, R., Zhang, Y., Zhang, L., Chen, M., Xin, L., & Zhang, L. (2025). The Effect of *Pseudomonas putida* on the Microbial Community in Casing Soil for the Cultivation of *Morchella sextelata*. Journal of Fungi, 11(11). 10.3390/jof11110775

