## Supplementary materials for "Effect of Mushroom-Bacteria Co-culture on Mushroom Growth and Antimicrobial Properties"

* Corresponding Author

**Supplementary Table S1**. Genes with mutations in the drug-resistant mutant of *B. subtilis* (BM2)

| **Categories** | ***Gene*** | **Description of the encoded protein** |
| --- | --- | --- |
| Sporulation | *skfC* | Killing non-sporulating siblings |
|  | *cwlH* | Autolysin, hydrolase, hydrolyzing cell wall |
|  | *sspB* | A small, acid-soluble spore protein protecting spore DNA |
| Quorum sensing | *srfAA and srfAB* | Synthesizing surfactin, a powerful biosurfactant that also facilitates biofilm formation. |
|  | *ppsABCD* | Synthesizing plipastatin, an antibiotic to kill fungus. |
| SPβ prophage: DNA repair /replication | *yobEH* | A TA module |
|  | *yokIJ* | A Type II restriction-modification system. YokI is the methyltransferase. YokJ is the restriction endonuclease. |
|  | *yoaM* | A putative specialized repair protein that prevents mutations by handling basic lesions |
|  | *uvrX* | Essential for error-free repair |
|  | *dutAB* | A two-gene module to ensure the SPβ genome remains intact and functional |
|  | *nrdF* | For DNA synthesis |
| SPβ prophage: TA modules | *yfjDE* | YfjD is the toxin targeting bacterial translation or replication machinery. YfjE is the antitoxin. |
|  | *yobEH* | YobH acts as an endoribonuclease or a translation inhibitor, targeting essential cellular processes. YobE is the antitoxin. |
|  | *yokFGH* | YokF (toxin) inhibits essential processes (typically DNA replication or protein synthesis); YokG the antitoxin is constantly produced to keep the toxin in check if SPβ exists. YokH is the chaperone stabilizing the toxin-antitoxin complex. The system ensures that SPβ is stable in the population. |
|  | *yobLK* | YobK is a potent ribonuclease (RNase). It cleaves cellular mRNA to shut down protein synthesis. |
|  | *yopCB* | YopC (toxin) is a putative NAD(+) phosphorylase, inhibiting essential metabolic processes to cause cell death. |
| PBSX prophage | *xlyA* | YylA is an endolysin, a cell wall hydrolase targeting the bacterial peptidoglycan. |
|  | *xkdP* | XkdP is part of the late operon for the tail of the PBSX prophage that is expressed only after the prophage has been induced (e.g., by DNA damage triggering the SOS response). |

**Supplementary Table S2.** Antimicrobial bioactive compounds from shiitake mushroom

| Compound(s) | Solubility | Antimicrobial properties | Reference |
| --- | --- | --- | --- |
| Lenthionine | Insoluble in water | Kills fungi and bacteria with more efficiency in killing fungi | Baral 2025  Avinash et al. 2016 |
| Lentinamycin | Insoluble in water | Antibacterial. Inhibits DNA replication. Found in the fermentation broth and mycelium. | Baral 2025 |
| Ergosterol | Insoluble in water | Antiviral and antibacterial. Abundant in organic solvent extract. | Sutthisa et al. 2025 |
| Carvacrol | Insoluble in water | Phenolic compound, disrupts microbial membranes | Baral 2025 |
| Copalic acid | Insoluble in water | Kills bacteria and inhibits biofilm formation | Baral 2025 |
| Cortinellin | Insoluble in water | Broad-spectrum antimicrobial activity by disrupting microbial membrane | Baral 2025 |
| Linoleic acid | Insoluble in water. Highly soluble in ethanol, DMSO and acetone. | More sensitive to gram-positive bacteria. It disrupts bacterial cell membrane and reduces biofilm formation | Yuyama et al. 2020 |
| Sesquiterpenes, steroids, anthraquinone | Insoluble in water. | Collectively, these are low molecular weight compounds having potent antimicrobial activity by inhibiting DNA , protein synthesis or cell division | Baral 2025;  Avinash et al. 2016 |
| Benzoic acid derivatives | Poorly soluble in acidic water. Soluble in neutral or basic water |  |  |
| Quinolones | poorly soluble at a neutral pH but solubility increases in strong acidic or basic conditions. |  |  |
| Erythritol | Soluble in water | Inhibit bacterial growth and biofilm formation | Baral 2025; de Cock et al. 2016; Lim et al. 2018 |
| Dianhydromannitol | Soluble in ethanol and water. | A major component (up to 20.1% or 21.8%) in the ethanol extracts of *A. bisporus* and *L.edodes*. | Erdogan 2022 |
| Lentin | Soluble in water. | Antifungal protein isolated from the fruiting body of mushrooms. | Baral 2025; Minutti et al. 2016; Muszyńska et al. 2017) |
| Lentinan | Soluble in water. | Polysaccharide. Reported to inhibit bacteria or fungi through its ability to improve immunity. | Baral 2025 |

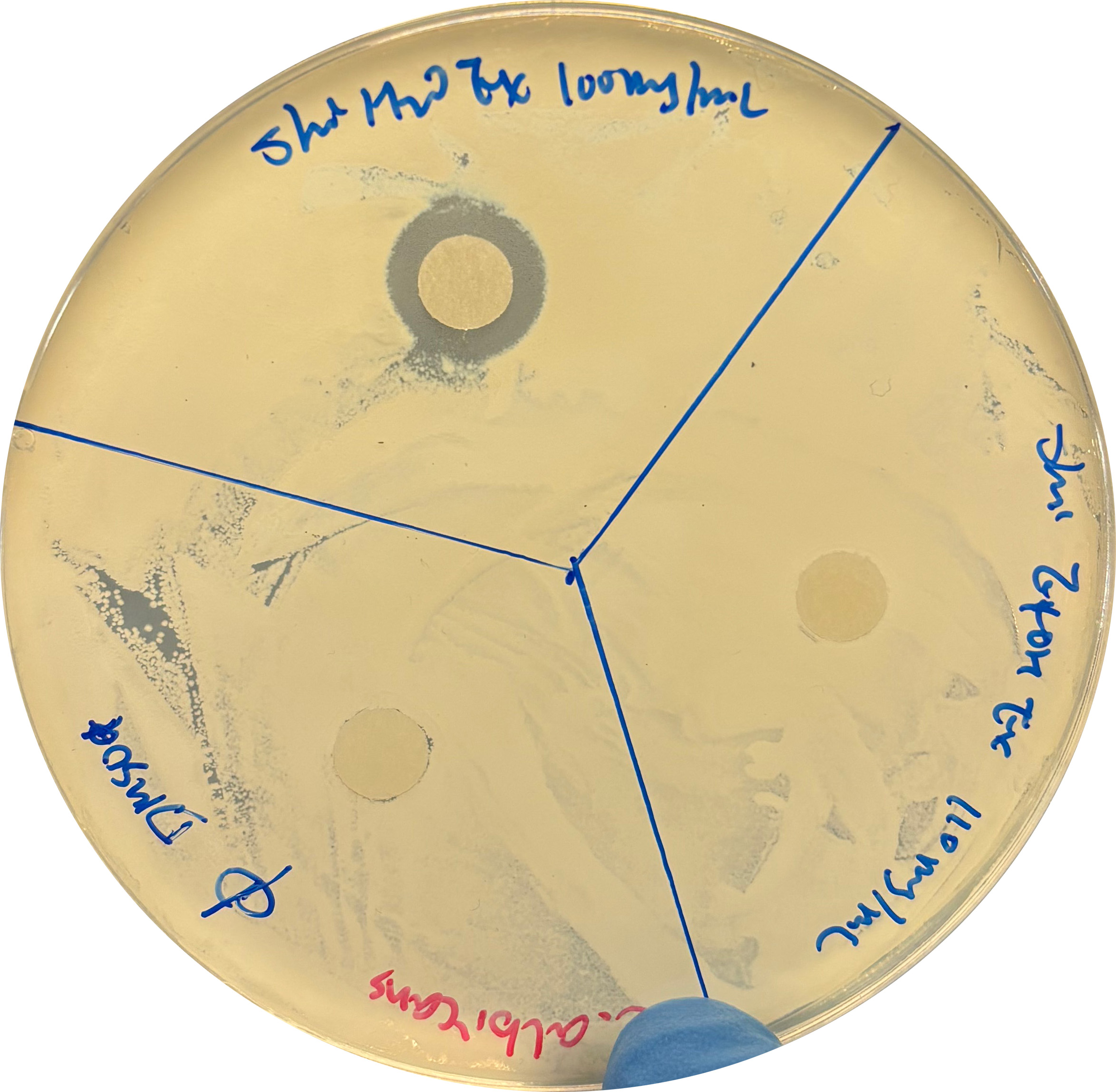

**Supplementary Fig. S1**. Shiitake water extract susceptibility diffusion disk test on *Candida albicans*. Top disk: shiitake water extract 100 mg/mL; left disk: DMSO, the solvent for ethanol extract; right disk: shiitake ethanol extract of 110 mg/mL.

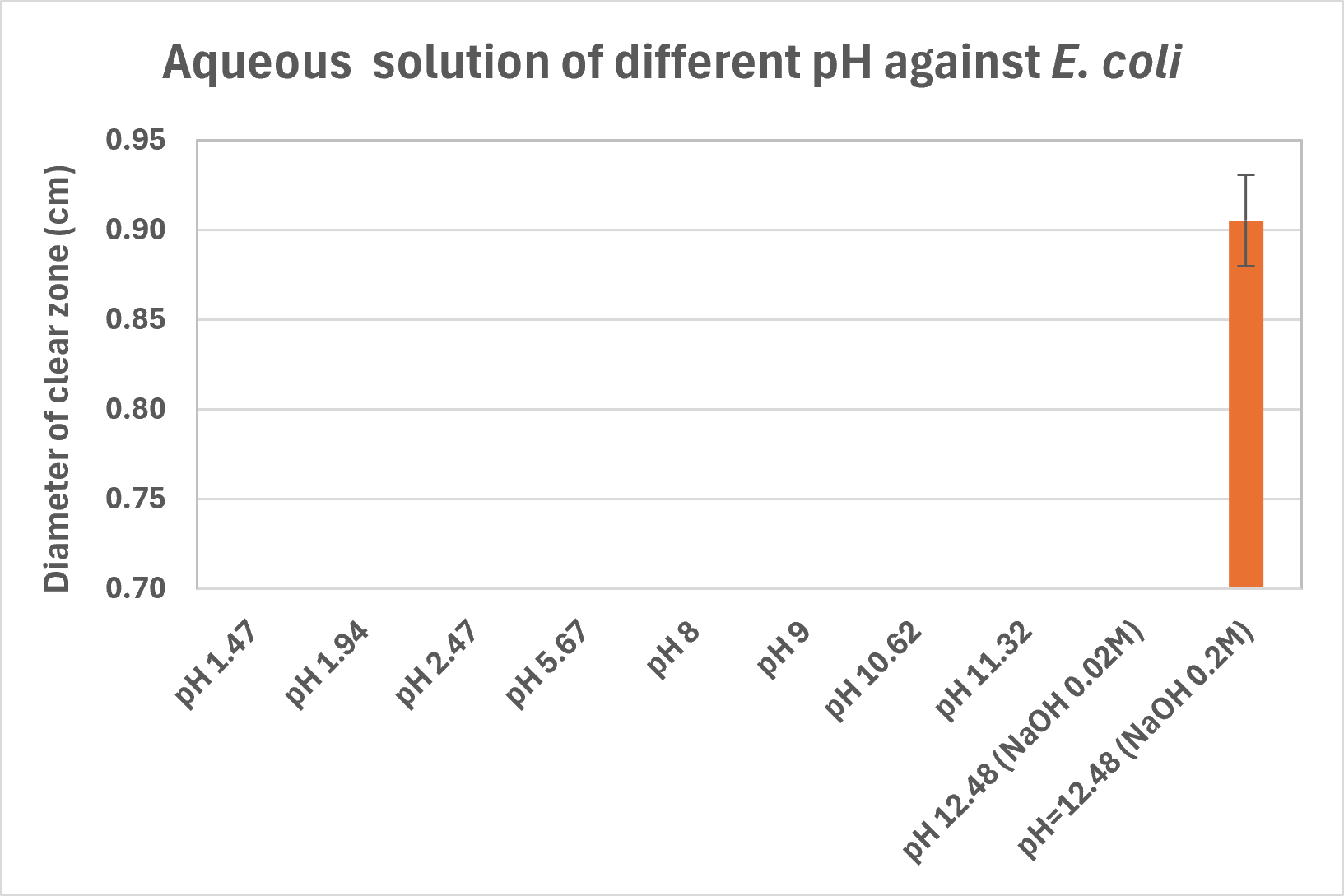

**Supplementary Fig.S2**. Susceptibility diffusion disk test of aqueous solutions of HCl and NaOH at different pH on the lawn of *E. coli*. Error bar: standard deviation (SD; n=3)

| 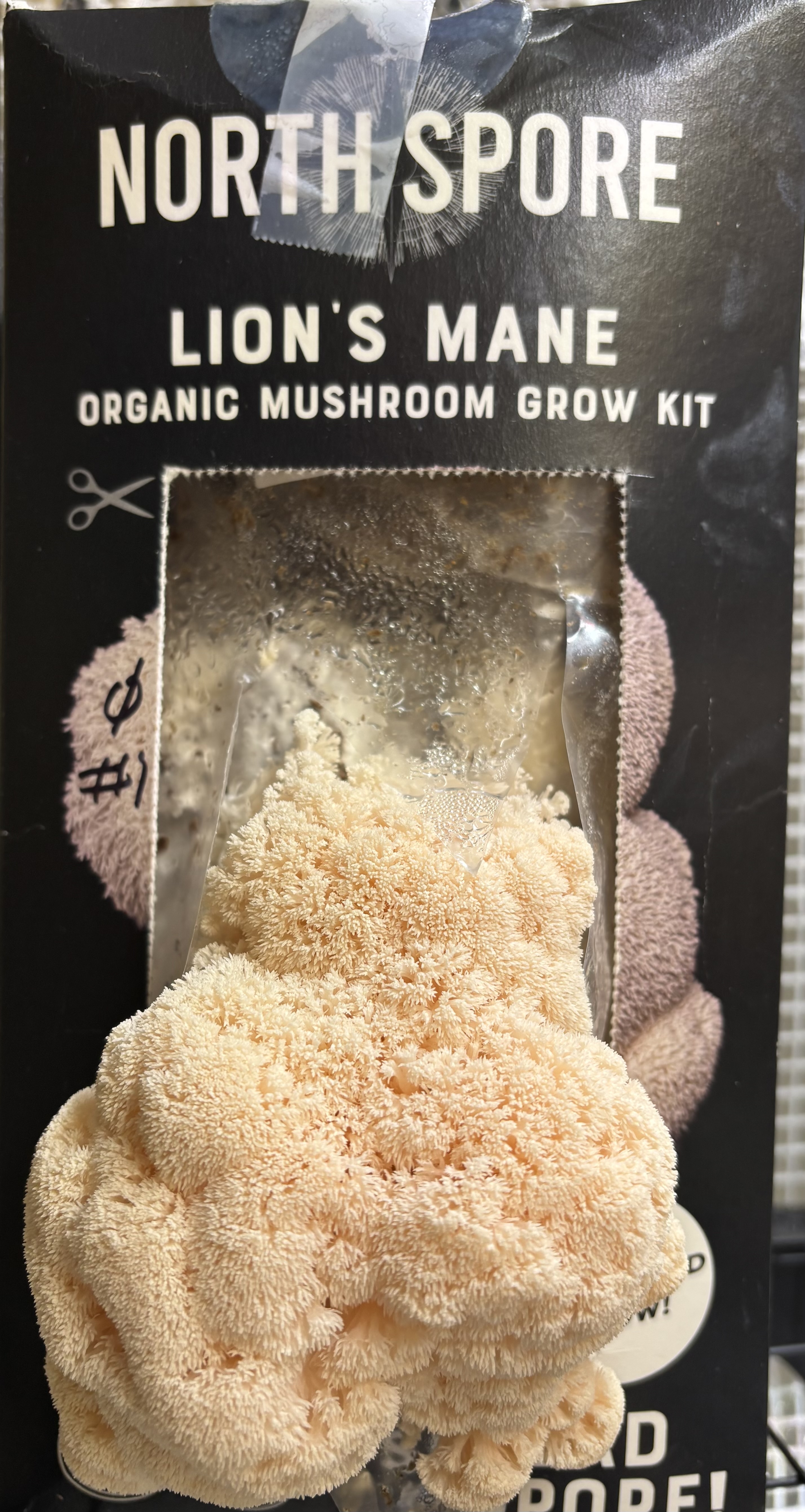 | 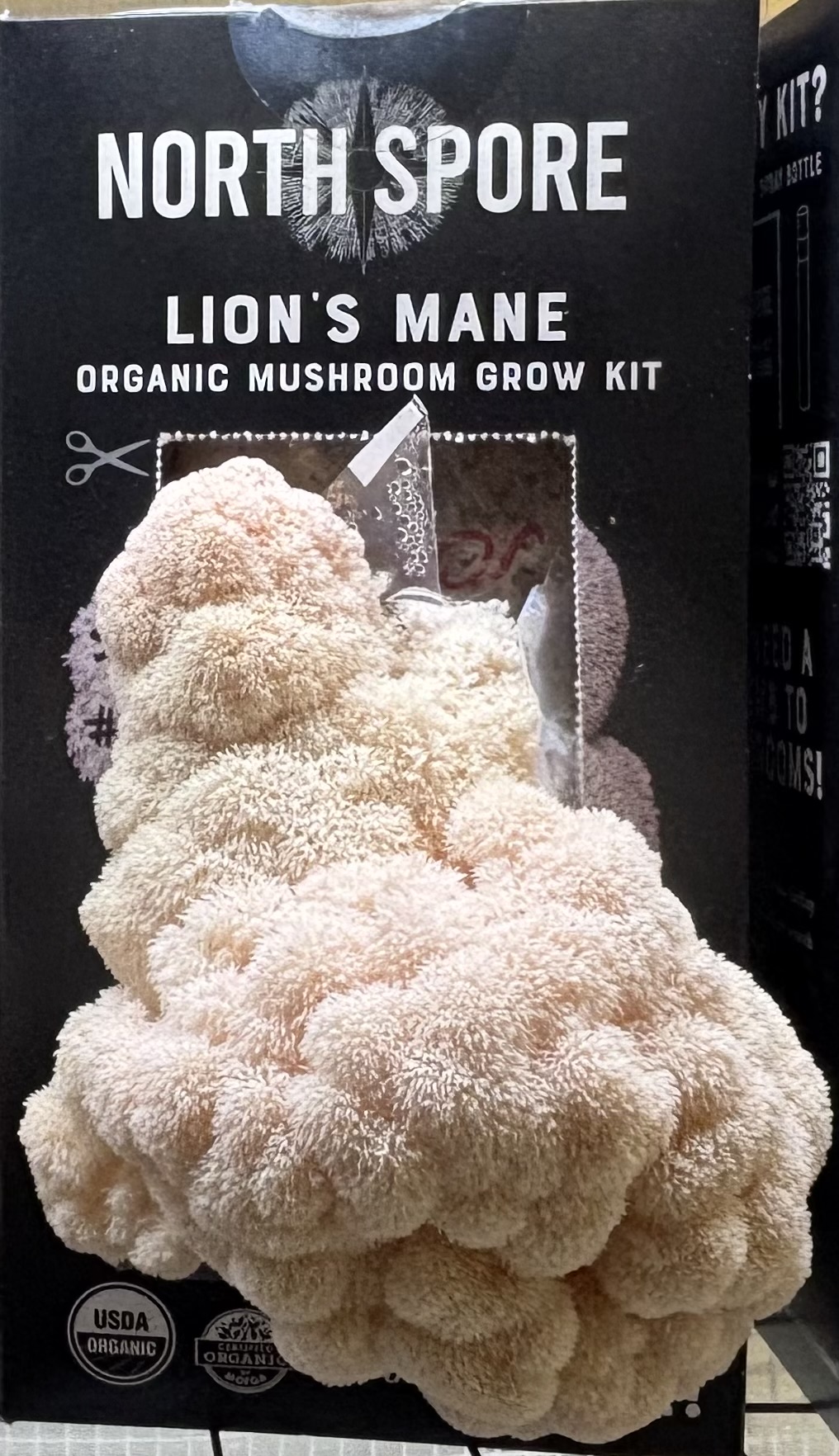 | 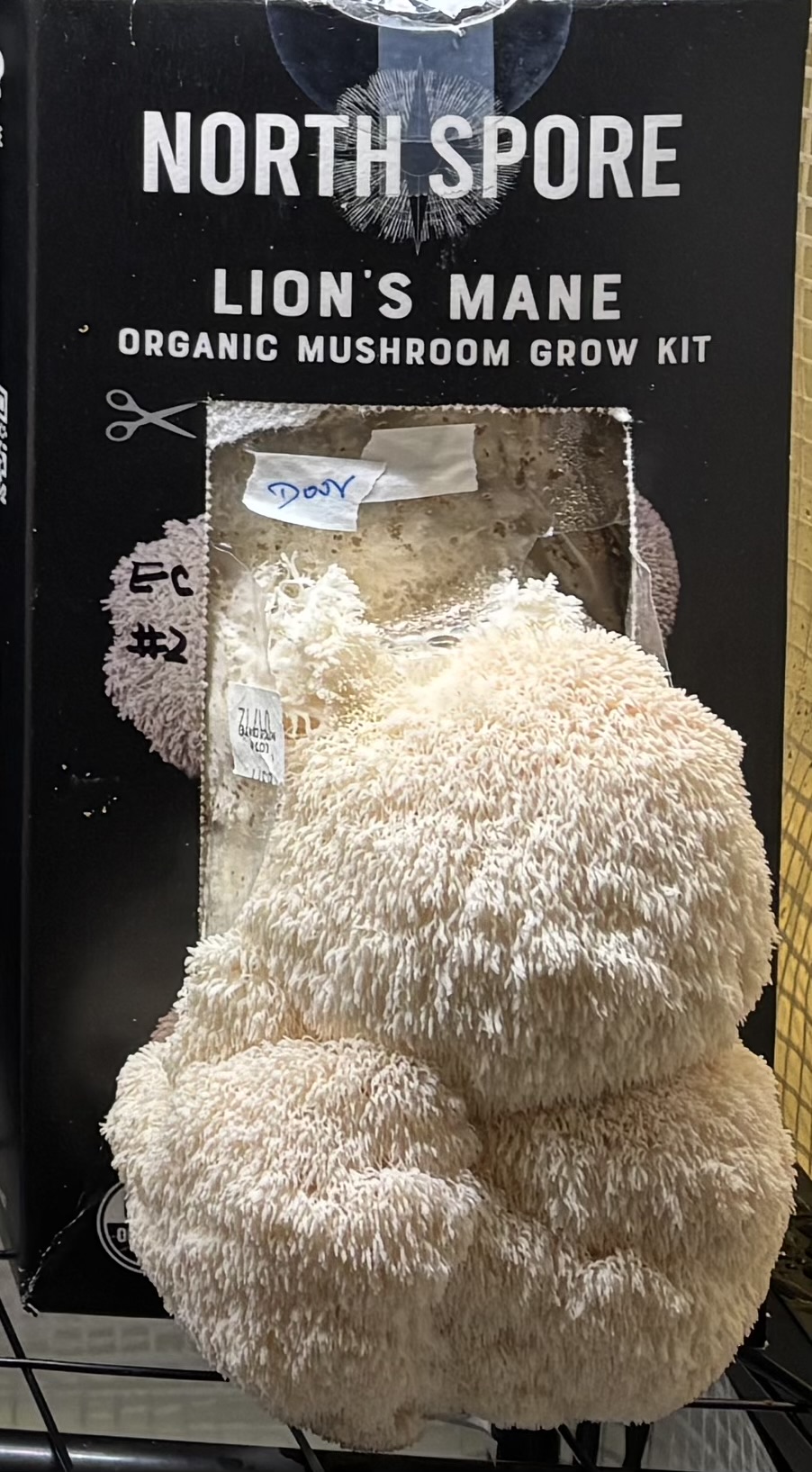 | 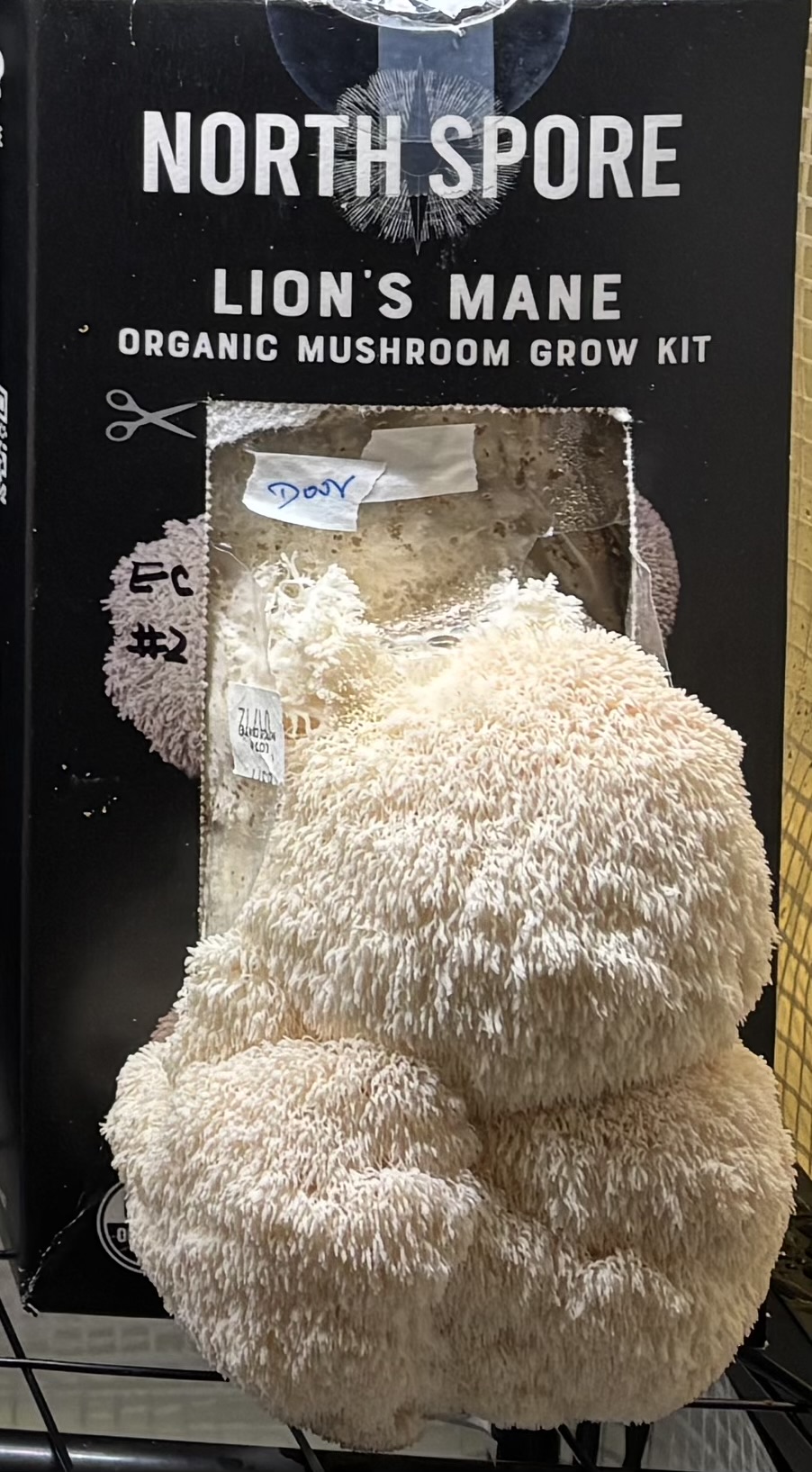 | 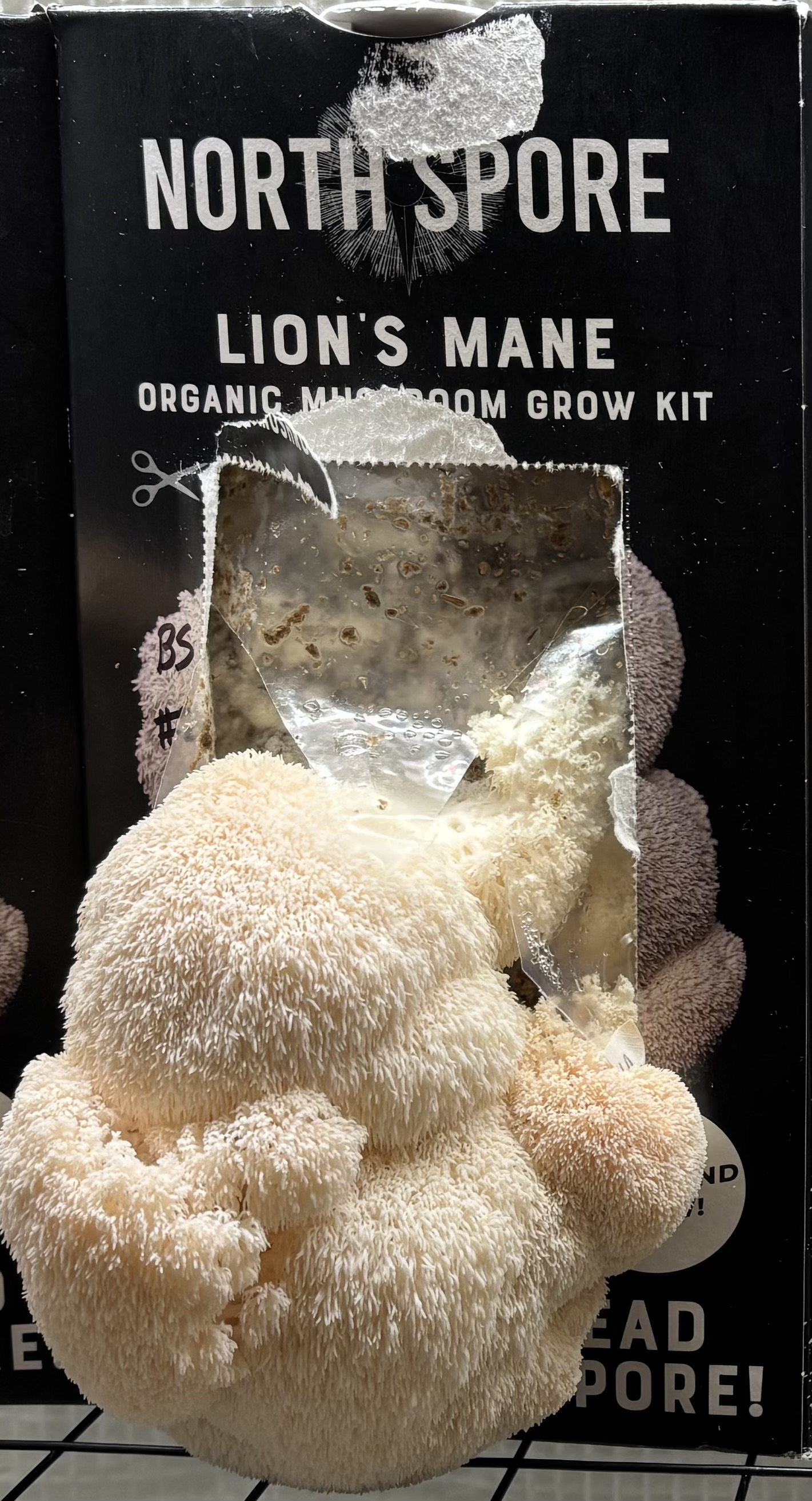 |
| --- | --- | --- | --- | --- |
| (A) | (B) | (C) | (D) | (E) |
| 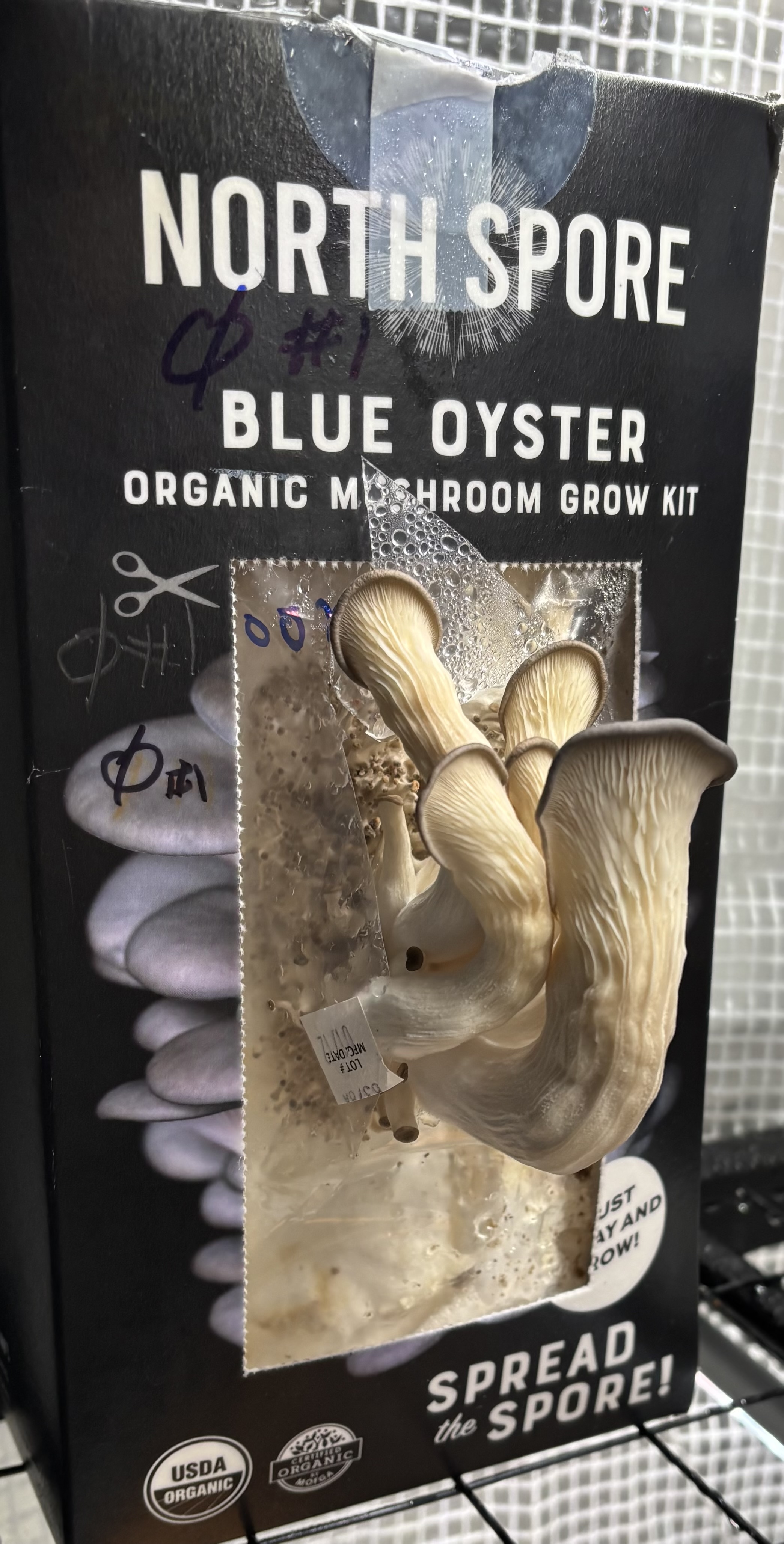 | 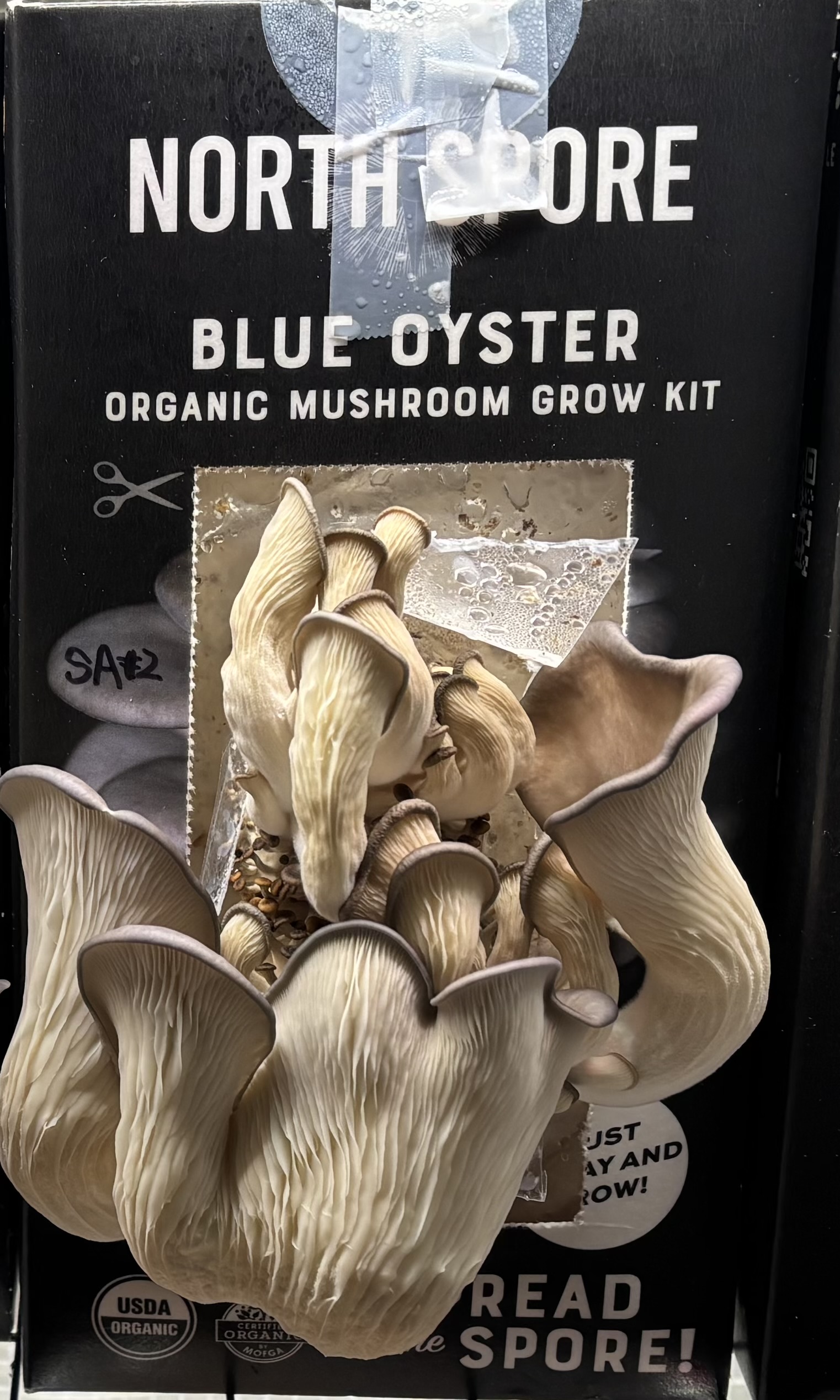 | 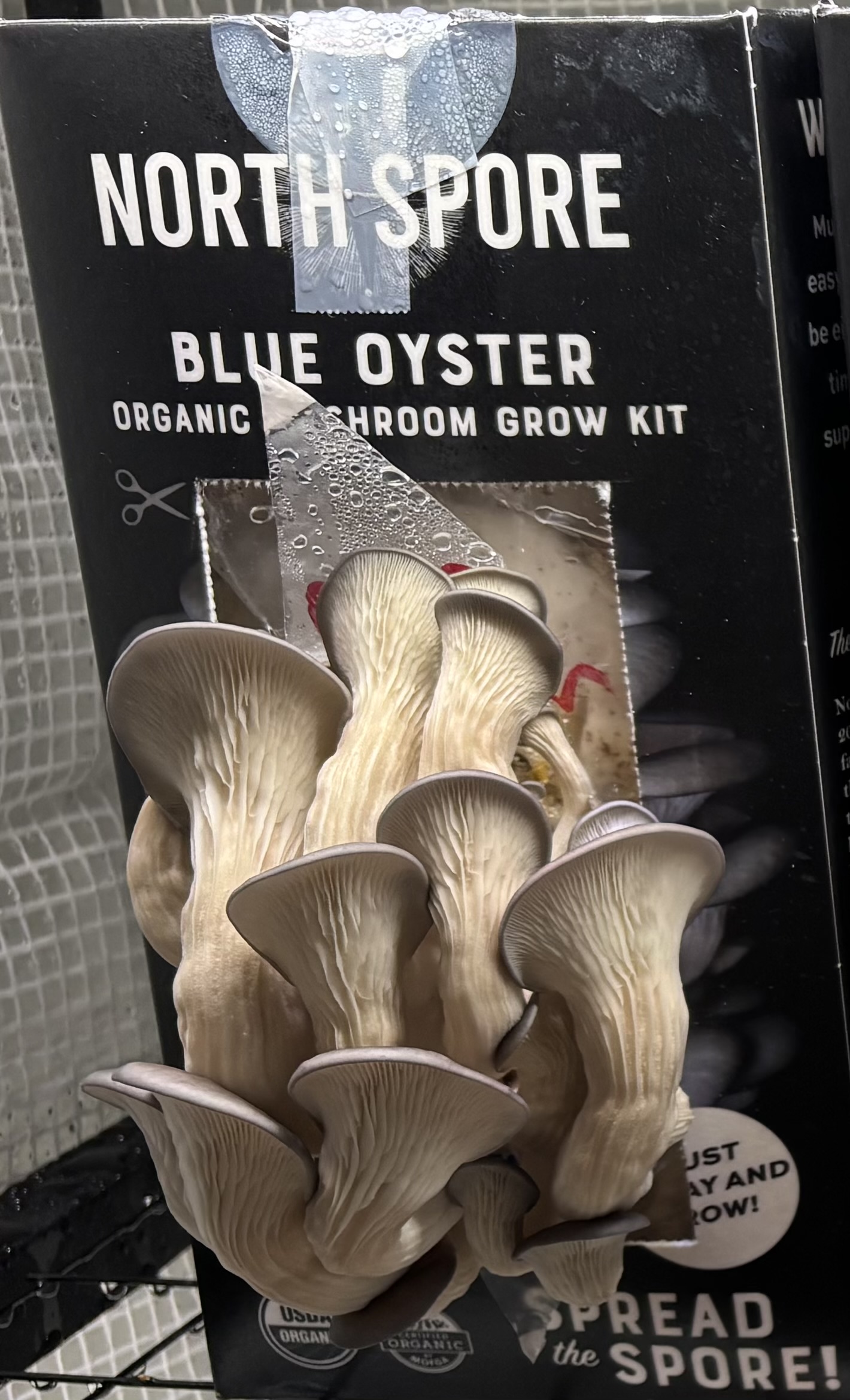 | 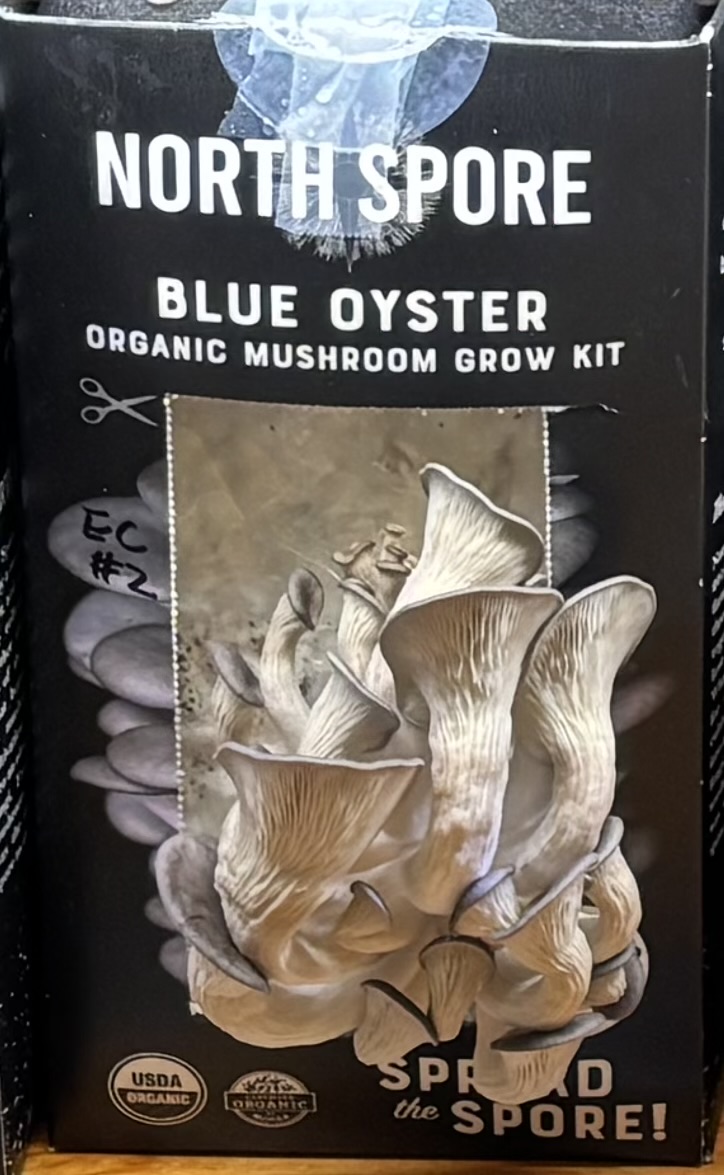 | 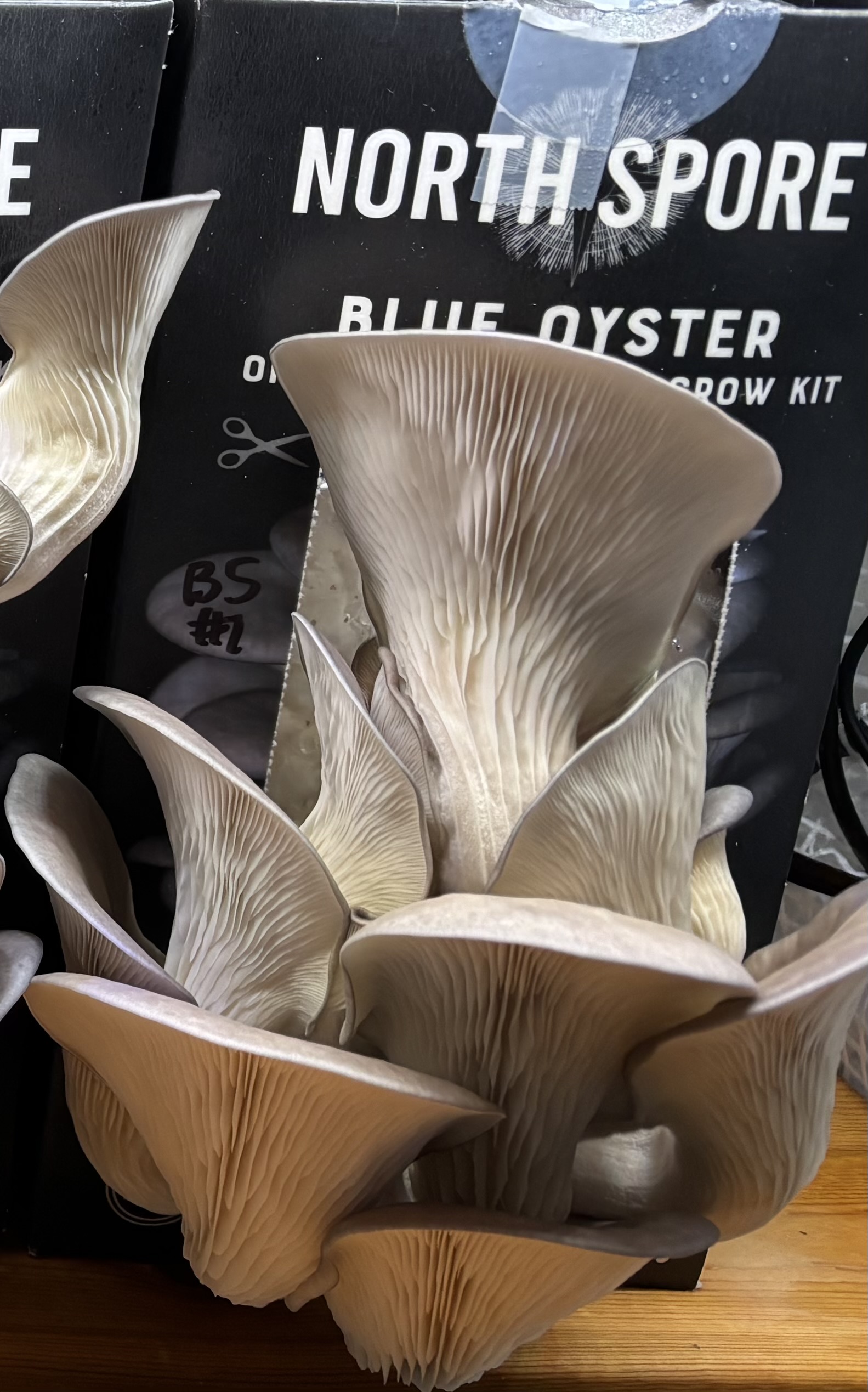 |
| (F) | (G) | (H) | (I) | (J) |

**Supplementary Fig.S3.** Top row: Co-culturing lion’s mane (LM) with (A) control; (B) *S. aureus*; (C) *P. aeruginosa*; (D) *E. coli* and (E) *B. subtilis* on Day 12. Bottom row: Co-culturing oyster mushroom (O) with (F) control; (G) *S. aureus*; (H) *P. aeruginosa*; (I) *E. coli* and (J) *B. Subtilis on Day 15.* The day 0 was when the substrate packages were sliced open. “Control” means water instead of bacteria being injected into mushroom substrate for co-culture.

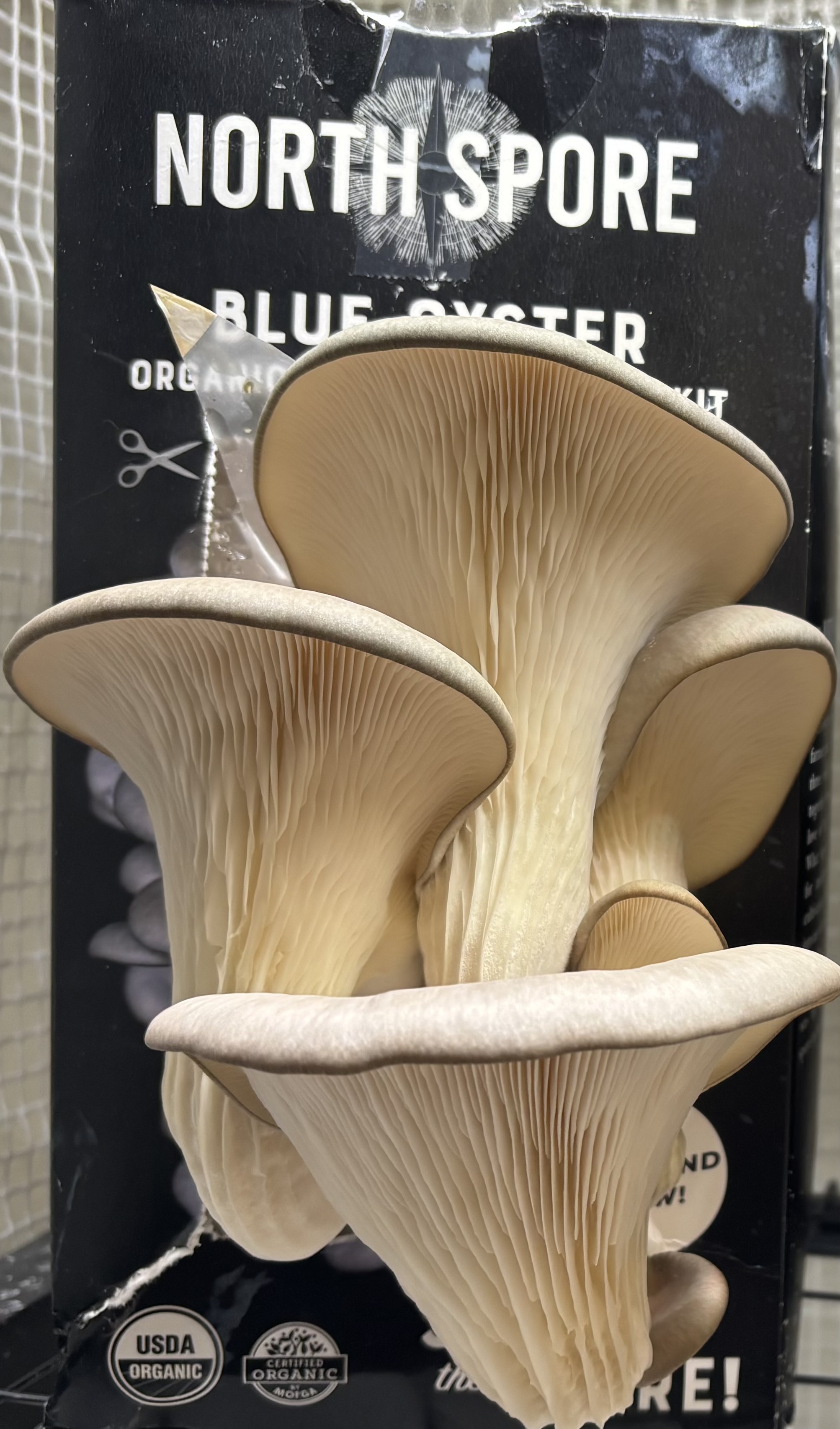

**Supplementary Fig. S4.** The oyster mushroom co-cultured with *P. aeruginosa* in the second flush.

**Supplementary References**

Avinash, J., Vinay, S., Jha, K., Das, D., Goutham, B. S., & Kumar, G. (2016). The unexplored anticaries potential of shiitake mushroom. *Pharmacognosy Reviews*, *10*(20), 100–104. <https://doi.org/10.4103/0973-7847.194039>

Baral, B. (2025). Holistic evaluation of shiitake mushrooms (*Lentinula edodes*): Unraveling its medicinal and therapeutic potentials. *Chemistry & Biodiversity*, *22*(12), e01244. <https://doi.org/10.1002/cbdv.202501244>

de Cock, P., Mäkinen, K., Honkala, E., Saag, M., Kennepohl, E., & Eapen, A. (2016). Erythritol is more effective than xylitol and sorbitol in managing oral health endpoints. *International Journal of Dentistry*, *2016*, 1–15. <https://doi.org/10.1155/2016/9868421>

Erdoğan Eliuz, E. A. (2021). Antibacterial activity and antibacterial mechanism of ethanol extracts of *Lentinula edodes* (shiitake) and *Agaricus bisporus* (button mushroom). *International Journal of Environmental Health Research*, 1–14. <https://doi.org/10.1080/09603123.2021.1919292>

Lim, J. H., Jeong, Y., Song, S.-H., Ahn, J.-H., Lee, J. R., & Lee, S.-M. (2018). Penetration of an antimicrobial zinc-sugar alcohol complex into *Streptococcus mutans* biofilms. *Scientific Reports*, *8*(1), 16154. <https://doi.org/10.1038/s41598-018-34366-y>

Minutti, L., Téllez-Téllez, M., R, D., S, T.-B., FJ, F., Santos-López, G., & Diaz-Godínez, G. (2016). Antimicrobial activity of a protein obtained from fruiting body of *Lentinula edodes* against *Escherichia coli* and *Staphylococcus aureus*. *Journal of Environmental Biology*, *37*(4), 619–623.

Muszyńska, B., Pazdur, P., Lazur, J., & Sułkowska-Ziaja, K. (2017). *Lentinula edodes* (shiitake) – biological activity. *Medicina Internacia Revuo*, *27*(108), 189–195. <https://interrev.com/mir/index.php/mir/article/view/17>

Sutthisa, W., Kamlangmak, P., & Srisawad, N. (2025). Antibacterial potential and chemical composition of shiitake mushroom ( *Lentinus edodes* (Berk.) Sing.) Extract against pathogenic bacteria. *Scientifica*, *2025*(1). <https://doi.org/10.1155/sci5/6089332>

Yuyama, K. T., Rohde, M., Molinari, G., Stadler, M., & Abraham, W.-R. (2020). Unsaturated fatty acids control biofilm formation of *Staphylococcus aureus* and other gram-positive bacteria. *Antibiotics (Basel, Switzerland)*, *9*(11), 788. <https://doi.org/10.3390/antibiotics9110788>
